# Metabolic strategies shared by basement residents of the Lost City hydrothermal field

**DOI:** 10.1101/2022.01.25.477282

**Authors:** William J. Brazelton, Julia M. McGonigle, Shahrzad Motamedi, H. Lizethe Pendleton, Katrina I. Twing, Briggs C. Miller, William J. Lowe, Alessandrina M. Hoffman, Cecilia A. Prator, Grayson L. Chadwick, Rika E. Anderson, Elaina Thomas, David A. Butterfield, Karmina A. Aquino, Gretchen L. Früh-Green, Matthew O. Schrenk, Susan Q. Lang

**Affiliations:** School of Biological Sciences, University of Utah, Salt Lake City, UT; Bigelow Laboratory for Ocean Sciences, 60 Bigelow Dr, East Boothbay, ME; Department of Molecular and Cell Biology, University of California, Berkeley, CA; Department of Biology, Carleton College, Northfield, MN; Joint Institute for the Study of Atmosphere and Ocean, University of Washington, Seattle, WA; Department of Earth Sciences, ETH Zurich, Zurich, Switzerland; Department of Earth and Environmental Sciences, Michigan State University, East Lansing, MI; School of the Earth, Ocean, and Environment, University of South Carolina, Columbia, SC, USA

## Abstract

Alkaline fluids venting from chimneys of the Lost City hydrothermal field flow from a potentially vast microbial habitat within the seafloor where energy and organic molecules are released by chemical reactions within rocks uplifted from Earth’s mantle. In this study, we investigated hydrothermal fluids venting from Lost City chimneys as windows into subseafloor environments where the products of geochemical reactions, such as hydrogen (H_2_), formate, and methane, may be the only available sources of energy for biological activity. Our deep sequencing of metagenomes and metatranscriptomes from these hydrothermal fluids revealed a few key species of archaea and bacteria that are likely to play critical roles in the subseafloor microbial ecosystem. We identified a population of *Thermodesulfovibrionales* (belonging to phylum *Nitrospirae*) as a prevalent sulfate-reducing bacterium that may be responsible for much of the consumption of H_2_ and sulfate in Lost City fluids. Metagenome-assembled genomes (MAGs) classified as *Methanosarcinaceae* and Candidatus Bipolaricaulota were also recovered from venting fluids and represent potential methanogenic and acetogenic members of the subseafloor ecosystem. These genomes share novel hydrogenases and formate dehydrogenase-like sequences that may be unique to hydrothermal and subsurface alkaline environments where hydrogen and formate are much more abundant than carbon dioxide. The results of this study include multiple examples of metabolic strategies that appear to be advantageous in hydrothermal and subsurface environments where energy and carbon are provided by geochemical reactions.

**IMPORTANCE:** The Lost City hydrothermal field is an iconic example of a microbial ecosystem fueled by energy and carbon from Earth’s mantle. Uplift of mantle rocks into the seafloor can trigger a process known as serpentinization that releases hydrogen and creates unusual environmental conditions where simple organic carbon molecules are more stable than dissolved inorganic carbon. This study provides an initial glimpse into the kinds of microbes that live deep within the seafloor where serpentinization takes place, by sampling hydrothermal fluids exiting from the Lost City chimneys. The metabolic strategies that these microbes appear to be using are also shared by microbes that inhabit other sites of serpentinization, including continental subsurface environments and natural springs. Therefore, the results of this study contribute to a broader, interdisciplinary effort to understand the general principles and mechanisms by which serpentinization-associated processes can support life on Earth and perhaps other worlds.

## INTRODUCTION

The fixation of carbon dioxide into organic carbon by autotrophic organisms is the foundation of all ecosystems on Earth. Even in subsurface environments, organic carbon is provided by fixation of carbon dioxide by chemoautotrophs or else from the degradation of organic carbon originally produced in photosynthetic ecosystems and transported into the subsurface. However, organic carbon can form abiotically in hydrothermal environments, particularly in those that favor a set of geochemical reactions collectively known as serpentinization (McCollom & Seewald, 2007; Martin et al., 2008). Microbial communities in serpentinizing environments are likely to benefit from the abiotic synthesis of simple organic compounds, but the processes and mechanisms that may allow this to occur are unknown.

The Lost City hydrothermal field is located near the summit of the Atlantis Massif, a submarine mountain formed by the uplift of ultramafic rocks from Earth’s upper mantle and emplacement onto the seafloor along a major fault zone (Kelley et al., 2005; Karson et al., 2006; Früh-Green et al., 2018). Serpentinization of the Atlantis Massif results in the generation of hydrogen gas (H_2_) and hydrothermal fluids that are rich in formate, methane, and perhaps other forms of organic carbon (Proskurowski et al., 2008; Lang et al., 2012, 2018). Dissolved inorganic carbon is vanishingly rare in the pH 9-11 hydrothermal fluids that vent from Lost City chimneys because it is either reduced to formate or methane or else precipitated as carbonate minerals (Proskurowski et al., 2008; Ternieten, Früh-Green & Bernasconi, 2021). Sulfate, in contrast, appears to be an available oxidant throughout the subseafloor because it is never completely consumed by the relatively moderate hydrothermal conditions within the Atlantis Massif (Kelley et al., 2005; Lang & Brazelton, 2020).

Dense biofilm communities coating the surfaces of Lost City chimneys are capable of utilizing this bounty of energy and carbon released from the mantle (Lang et al., 2018; McGonigle, Lang & Brazelton, 2020). However, these biofilms form in mixing zones where warm, anoxic hydrothermal fluids vent into cold, oxic seawater. These conditions may not be representative of subseafloor environments within the Atlantis Massif where habitats are probably confined to sparsely distributed fractures and channels within rocks that have limited exposure to seawater (Früh-Green et al., 2018; Motamedi et al., 2020). In particular, dissolved inorganic carbon is provided by ambient seawater to chimney biofilm communities, while its availability is severely limited in subseafloor habitats dominated by the products of serpentinization.

The microbiology of fluids venting from Lost City chimneys has been explored in only one study (Brazelton et al., 2006), as all other microbiological research at Lost City has focused on the chimney biofilms (Brazelton et al., 2010, 2011; Lang et al., 2012, 2018; McGonigle, Lang & Brazelton, 2020; Lang & Brazelton, 2020). That early census of microbial diversity identified several novel 16S rRNA sequences, but they were poorly classified due to the limitations of microbial taxonomy at the time (Brazelton et al., 2006). In particular, the presence of potential sulfate-reducing bacteria (SRB) in Lost City fluids has been a mystery despite clear biogeochemical trends that indicate widespread SRB activity in the subseafloor (Lang et al., 2018; Lang & Brazelton, 2020).

A deep-sea expedition to the Lost City in 2018 was designed to fill this knowledge gap by investigating the microbiology and biogeochemistry of fluids venting from Lost City chimneys (Lang et al., 2021). We exploited natural biogeochemical trends in fluids venting from distinct chimney locations within the Lost City field to test hypotheses about subseafloor microbial metabolic activity. Here we report initial results from the sequencing of DNA and RNA in Lost City fluids, including the first sequences of metagenomes and metatranscriptomes from Lost City hydrothermal fluids. We identify a few key archaea and bacteria that appear to be indicative of subseafloor habitats strongly influenced by serpentinization. These results highlight metabolic strategies and adaptations that are common to life fueled by the products of serpentinization, including the potential use of formate and other simple forms of organic carbon as the primary sources of carbon for the ecosystem.

## RESULTS

### Characteristics of Lost City hydrothermal fluid samples

Hydrothermal fluid samples were collected from actively venting chimneys at the Lost City hydrothermal field (**Figure 1; Supplemental Figure S1**) using ROV *Jason* during the 2018 Lost City expedition aboard R/V *Atlantis* (AT42-01). This study includes 39 samples of hydrothermal fluids that were dedicated to DNA and RNA sequencing, including analyses of amplicon sequence variants (ASVs), metagenomes, and metatranscriptomes (**Table 1**; **Supplemental Table S1).**

**Figure 1.**
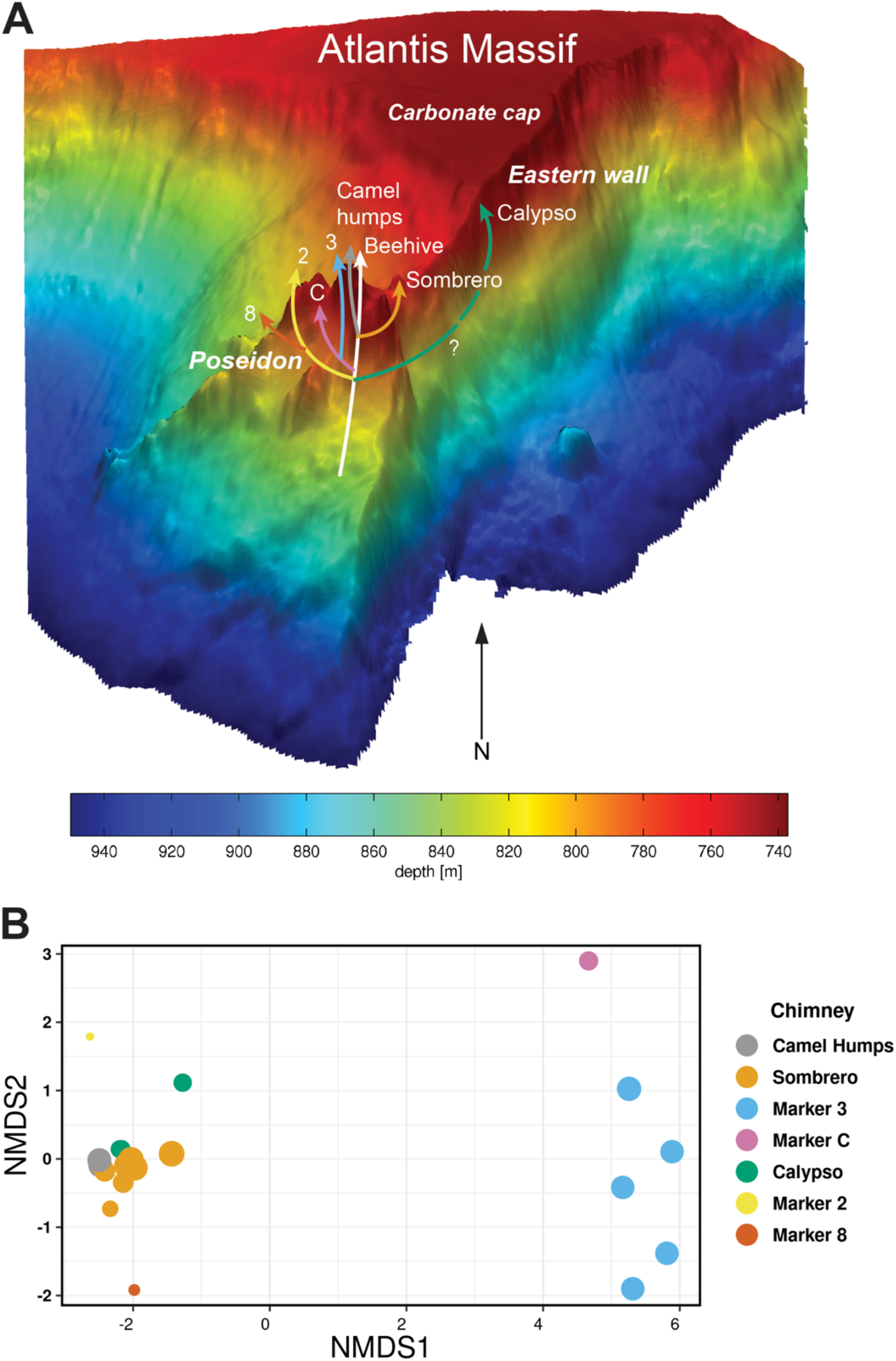
The Lost City hydrothermal field is located at 30°N, west of the Mid-Atlantic Ridge, on the southern wall of the Atlantis Massif. Part A shows a three-dimensional view of the field (after Kelley et al., 2005) featuring the massive Poseidon structure, which is composed of several actively venting chimneys. Hypothetical flow paths are informed by the aqueous geochemistry results reported here, by Aquino et al. (in review), and by prior studies referenced in the main text. Part B is a non-metric multidimensional scaling (NMDS) ordination of 16S rRNA amplicon sequence data where each data point represents the microbial community composition of one hydrothermal fluid sample. Sizes of data points are scaled to the measured sulfate concentration of that sample (Table 1).

**Table 1.** Overview of hydrothermal fluid samples collected from Lost City chimneys. Cell numbers are the median of all samples collected from that location. Temperatures and chemistry values are reported for one representative sample collected from that location, typically the sample for which the most chemistry and/or sequence data was available.

| *Chimney location* | *Meta-genome libraries* | *Meta-transcriptome libraries* | *16S rRNA amplicon libraries (DNA)* | *16S rRNA amplicon libraries (RNA)* | *Cells per mL* | *Max T (°C)* | *pH* | *Sulfide (mmol/kg)* | *Sulfate (mmol/kg)* | *Mg (mmol/kg)* |
| --- | --- | --- | --- | --- | --- | --- | --- | --- | --- | --- |
| *Camel Humps* | 2 | 0 | 4 | 0 | 4.3 x 10<sup>4</sup> | 41 | 8.1 | 0.02 | 24 | 47 |
| *Sombrero1* | 3 | 1 | 7 | 1 | 5.9 x 10<sup>4</sup> | 13 | 8.0 | 0.002 | 27 | 54 |
| *Sombrero2* | 2 | 0 | 6 | 1 | 6.3 x 10<sup>4</sup> | 52 | 8.7 | 0.12 | 18 | 36 |
| *Marker 3* | 2 | 0 | 6 | 0 | 6.3 x 10<sup>4</sup> | 21 | 8.6 | 0.20 | 23 | 45 |
| *Marker C* | 0 | 0 | 1 | 1 | 3.8 x 10<sup>4</sup> | 80 | 10.1 | 0.39 | 16 | 31 |
| *Calypso* | 2 | 0 | 6 | 1 | 4.8 x 10<sup>4</sup> | 24 | 9.3 | 1.3 | 15 | 30 |
| *Marker 2* | 2 | 1 | 3 | 1 | 2.6 x 10<sup>4</sup> | 58 | 9.5 | 1.8 | 8 | 15 |
| *Marker 8* | 0 | 0 | 1 | 0 | 7.5 x 10<sup>4</sup> | 42 | 9.9 | 1.8 | 9 | 19 |

The fluid samples ranged from those that were barely distinguishable from ambient seawater (∼11 °C, pH 8) to warm and highly alkaline hydrothermal fluids (∼80 °C, pH 10). Direct counts of visible cells showed little variability among fluids, with densities approximately 2-8 × 10^4^ mL^-1^ in all samples, although the two samples with the highest temperatures had the least number of cells (**Table 1**).

Fluids venting from Markers 3 and C contained ASV compositions that were notably distinct from those of all other fluids (**Figure 1**), including high relative abundances of *Thermodesulfovibrionia*, *Desulfotomaculum*, and *Bipolaricaulota* (**Figure 2**; Supplemental Table S2). In addition, Marker 3 fluids were rich in metagenomic sequences classified as family *Methanosarcinaceae,* which includes the dominant archaeal phylotype previously detected in Lost City chimneys (Schrenk et al., 2004; Brazelton et al., 2010, 2011). The greater representation of archaeal sequences in the metagenomes suggests a bias against archaeal sequences in the ASV dataset.

**Figure 2.**
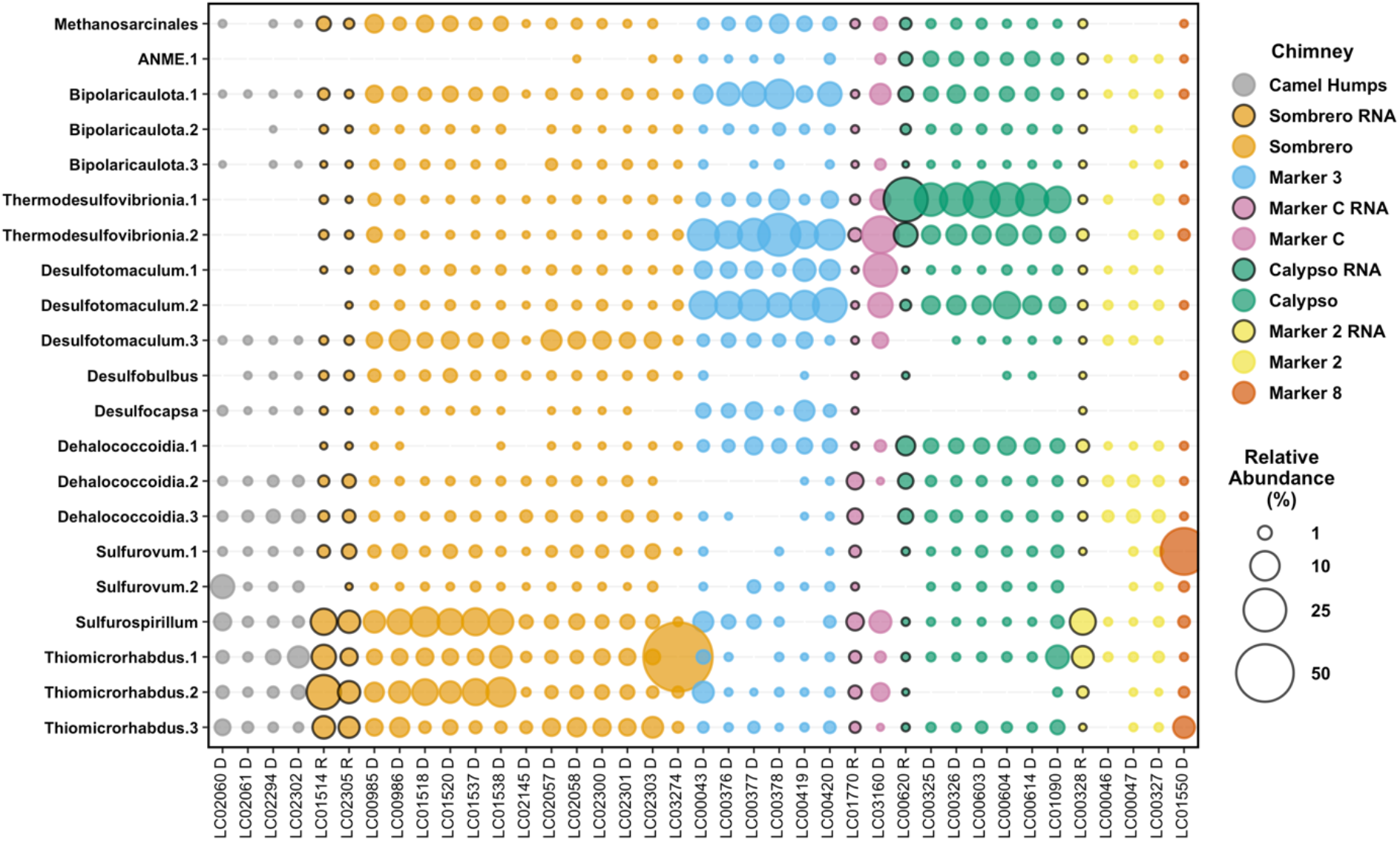
Relative abundances of selected ASVs in Lost City hydrothermal fluid samples. Amplicon libraries were generated from both DNA and RNA extractions; bubbles representing relative abundances in RNA libraries are highlighted with black borders. ASVs were selected to highlight the taxa that were the focus of this study, as well as additional taxa that are expected to be associated with hydrothermal environments and provide context for interpreting differences among fluid samples. A full table of ASV counts is provided in **Supplemental Table S2**.

Fluids venting from Camel Humps contained a remarkably even distribution of ASVs that included *Sulfurovum*, *Sulfurospirillum*, and *Thiomicrorhabdus* at similar abundances as taxa typically associated with ambient seawater (e.g., *Alteromonas*, *Roseobacter, Halomonas*). The overall microbial community structure of Sombrero fluids is broadly similar to that of Camel Humps fluids, although warmer and more sulfidic Sombrero fluids included greater proportions of taxa that were also abundant in fluids from Markers 3 and C (**Figure 2**). Fluid samples from the chimneys at Markers 2 and 8 were dominated by bacteria that are ubiquitous in chimney surface biofilm communities (Brazelton et al., 2006, 2010).

In general, the proportion of ambient seawater in each hydrothermal fluid sample, as measured by Mg concentration, did not predict the presence of microbes likely to inhabit anoxic, subseafloor environments. Instead, the distribution of anaerobic taxa most likely to be strongly linked with serpentinization (e.g., *Methanosarcinaceae, Thermodesulfovibrionia*, *Desulfotomaculum*, and *Bipolaricaulota*) was strongly chimney-specific, indicating a strong influence of subsurface conditions that is only weakly mitigated by the mixture of seawater during sampling. Detailed comparisons of the hydrothermal fluid samples are provided in the Supplemental Material.

### Metagenome-Assembled Genomes (MAGs)

A total of 305 MAGs with at least 50% estimated completion were recovered from the pooled “all fluids” assembly and the six chimney-specific assemblies (**Supplemental Figure S3**; **Supplemental Table S4**). MAGs that were representative of the taxa enriched in Markers 3 and C, as well as MAGs that contained key genes associated with the metabolism of H_2_, sulfate, formate, and methane, were selected for additional analyses.

Re-assembly and manual refinement of these sequences (**Supplemental Material**) resulted in 30 refined and curated MAGs (**Figure 3**) that are at least medium-quality (>50% complete, <10% redundancy, (Bowers et al., 2017). Generally, these MAGs are most abundant in Marker 3, Calypso, or Sombrero, and they are nearly absent in Camel Humps and Marker 2. (Unfortunately, metagenomic sequences could not be obtained from Marker C or Marker 8). A single *Methanosarcinaceae* MAG was especially abundant in the fluids from Marker 3 (**Figure 3**). Each of these MAGs represent new species (or else represent novel taxa we have previously detected at Lost City), and our phylogenetic analyses indicate that the *Methanosarcinaceae*, *Thermodesulfovibrionales*, and Bipolaricaulota MAGs most likely represent novel genera (**Supplemental Figures S5-S7**).

**Figure 3.**
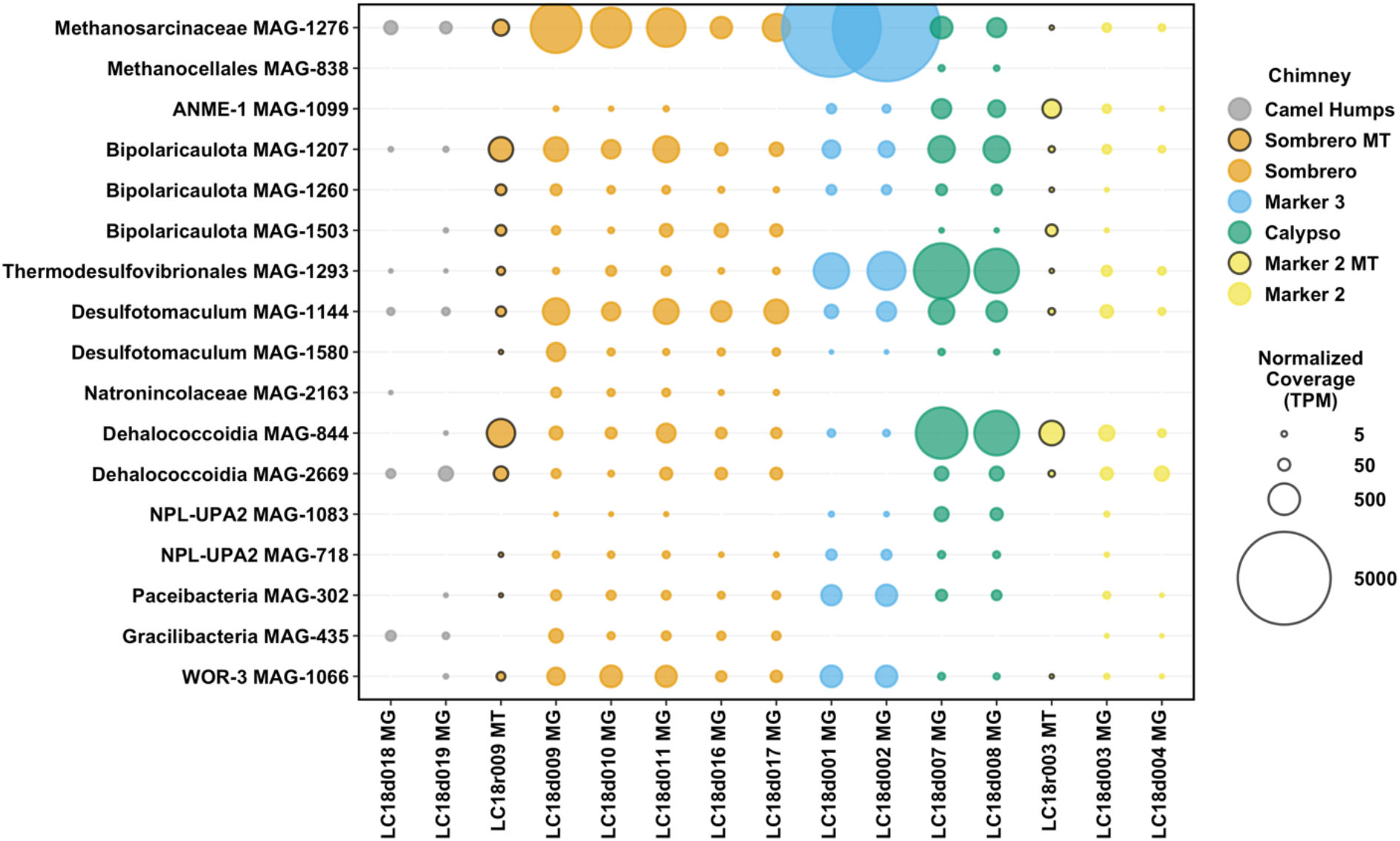
Abundance of refined MAGs in Lost City hydrothermal fluid samples. Total mapped read coverage was normalized to genome size and to the size of the metagenome or metatranscriptome library. The final normalized coverage is reported as a proportional unit (transcripts/fragments per million; TPM) suitable for cross-sample comparisons. Bubbles representing coverage in metatranscriptomes (MT), rather than metagenomes (MG), are highlighted with black borders. For clarity, not all MAGs are shown; a full coverage table is provided in **Supplemental Table S4**.

Below, we briefly describe key features of these MAGs that seem relevant to an initial exploration of the Lost City subseafloor ecosystem, focusing on genes associated with the metabolism of H_2_, formate, sulfur, and methane. Additional information about each MAG is reported in the **Supplemental Material**, including detailed descriptions of genomic content and predicted protein functions (**Supplemental Tables S5-S6)**.

### Hydrogenases

[NiFe]-hydrogenases typically associated with H_2_ oxidation were found in *Thermodesulfovibrionales* MAG-1293 (HyaAB), *Methanocellales* MAG-838 (HyaAB), and Bipolaricaulota MAG-1503 (HoxYH) (**Figure 4**). Of these, the *Thermodesulfovibrionales* MAG was by far the most abundant in venting fluids (**Figure 3**). *Methanosarcinaceae* MAG-1276 encodes two hydrogenases associated with methanogenesis: F_420_-reducing hydrogenase (FrhAB) and Ech hydrogenase (EchCE). It also encodes a formate dehydrogenase that can provide electrons to MvhD and HdrABC instead of the H_2_-oxidizing Vho/Vht enzyme (Costa et al., 2013). Thus, Lost City *Methanosarcinaceae* may power methanogenesis with electrons from both H_2_ and formate. The same MvhD-HdrABC complex, without FDH, was also found in MAGs classified as ANME-1, *Natronincolaceae*, and Bipolaricaulota (**Supplemental Table S5**).

**Figure 4.**
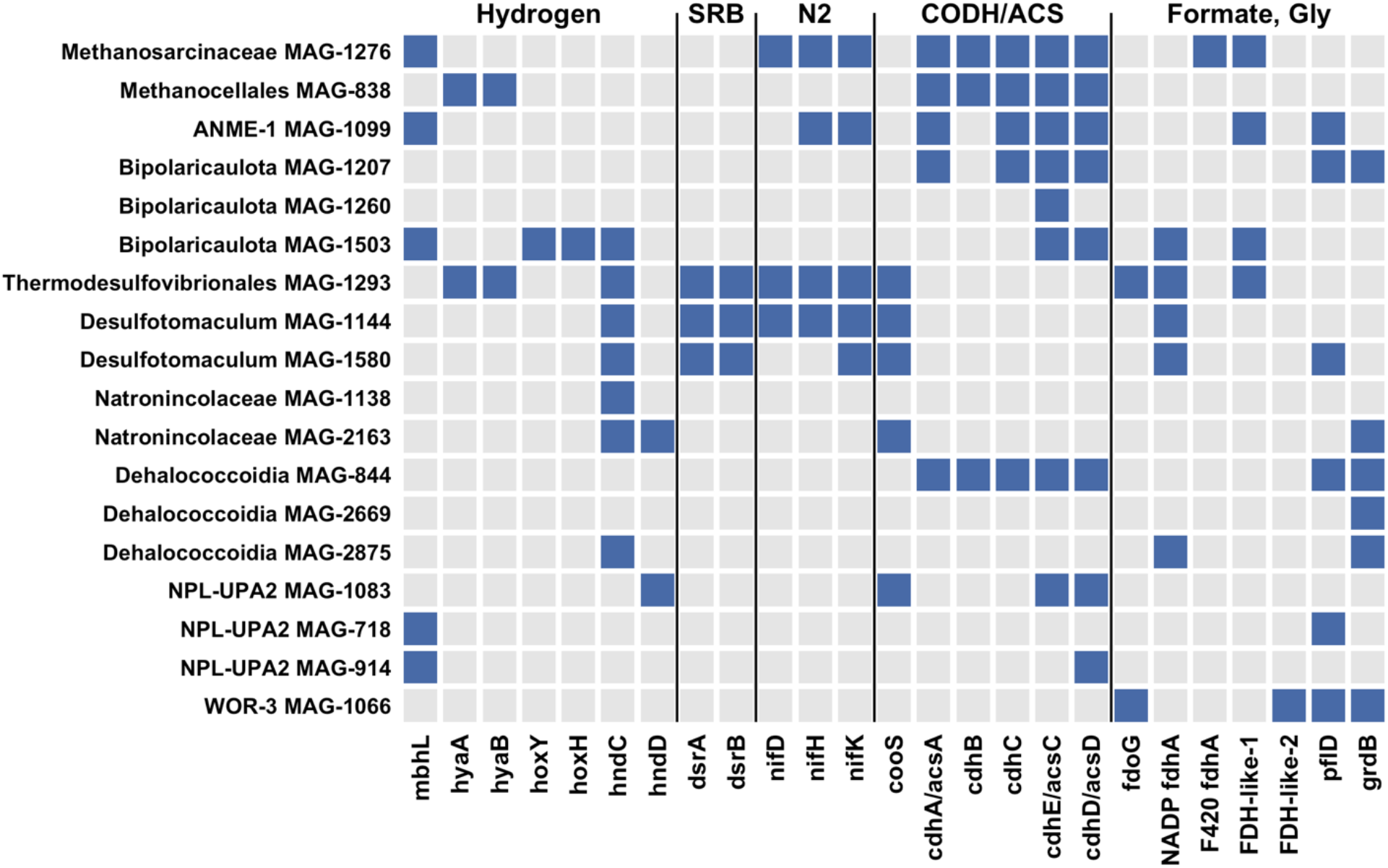
Presence and absence of key genes in refined MAGs. Genes defined by KEGG Orthology (see **Supplemental Table S5**) were selected to highlight potential metabolic capabilities to metabolize hydrogen gas, to reduce sulfate to sulfide (SRB), to fix nitrogen (N_2_) gas, to fix carbon dioxide via the Wood-Ljungdahl pathway (CODH/ACS), and to utilize formate or glycine as carbon sources. Patescibacteria MAGs (including Paceibacteria and Gracilibacteria) are not shown here because they lack all of the gene shown here.

In addition, the *Methanosarcinaceae* and ANME-1 MAGs contain a complete 14-gene cluster (mbhA-N) encoding membrane-bound hydrogenase (Mbh) (**Figure 5; Supplemental Figure S8**). For each predicted gene in the cluster, the homologs in the *Methanosarcinaceae* and ANME-1 MAGs are more similar to each other than to any other sequences in public databases. The same gene cluster, with conserved synteny, is also found in methanogens belonging to the order *Methanomicrobiales* and in heterotrophs of the order *Thermococcales* (Thauer et al., 2010). The MbhL subunits from these methanogens have only 42-45% identities with the Lost City MbhL sequences reported here, which have greater similarly (∼49% identities) to MbhL sequences from *Thermococcus*. Bipolaricaulota MAG-1503 also includes a predicted MbhL sequence, which is most closely related to two Bipolaricaulota MAGs from hydrothermal systems: the Mid-Cayman Rise (Zhou et al., 2020) and Guaymas Basin (Dombrowski, Teske & Baker, 2018) (**Figure 5**).

**Figure 5.**
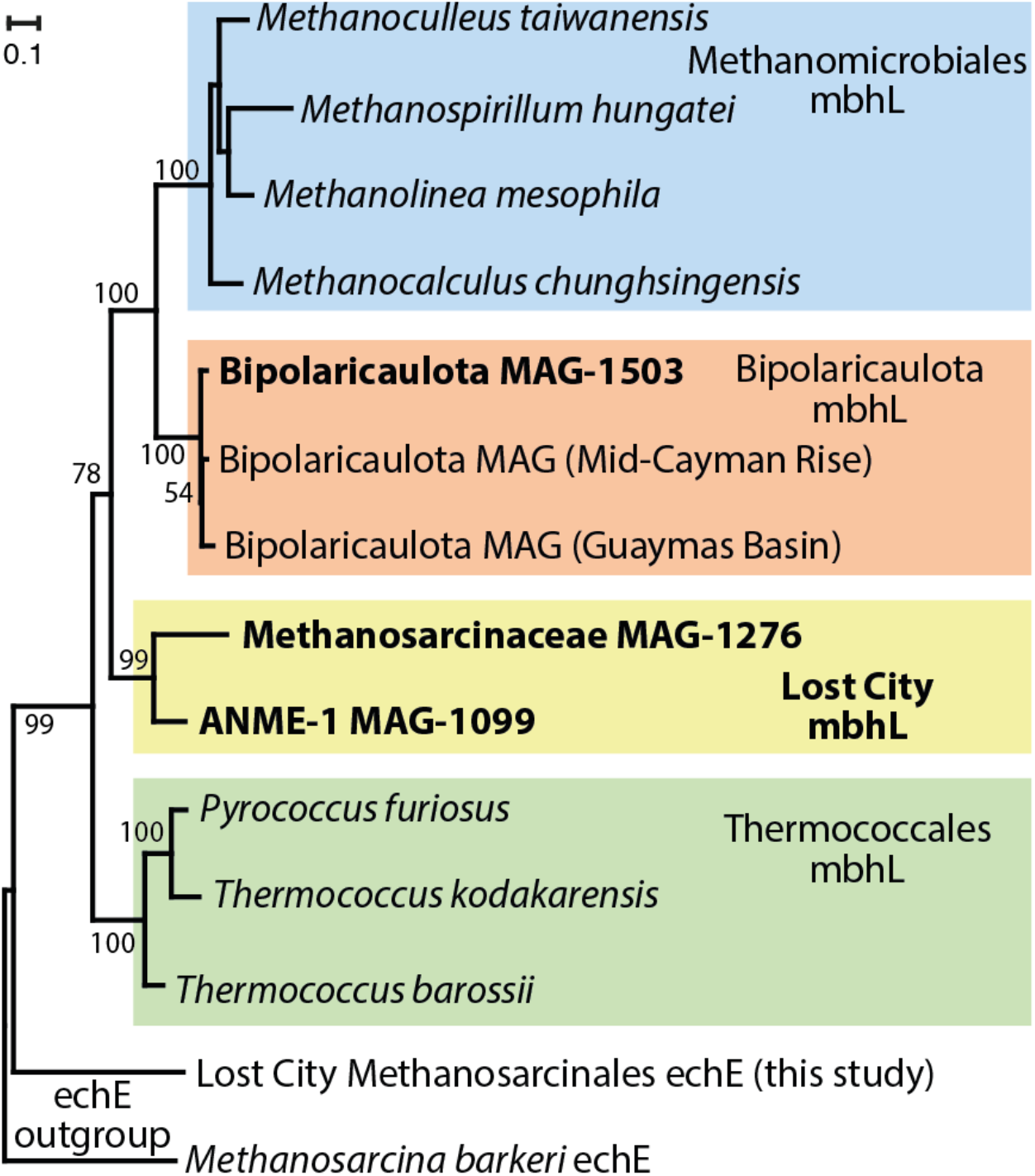
Phylogeny of the large catalytic subunit of membrane-bound hydrogenase (mbhL). Sequences identified in refined MAGs from this study are highlighted in bold font. The two archaeal sequences from Lost City (*Methanosarcinaceae* and ANME-1) form their own clade apart from all known mbhL sequences. The mbhL sequence from a Lost City Bipolaricaulota MAG clusters together with Bipolaricaulota MAGs from other hydrothermal environments. Bootstrap support values are shown for each node. An expanded version of this figure including the gene order for the mbh gene cluster is provided as **Supplemental Figure S4**.

[NiFe]-hydrogenase sequences (HyaAB) were also highly abundant in Sombrero and Camel Humps fluids (**Supplemental Table S7**), where they were primarily encoded by *Thiomicrorhabdus*. We did not prioritize the analysis of *Thiomicrorhabdus* MAGs because our prior work indicated that they inhabit oxygenated biofilm communities on chimney surfaces (Brazelton & Baross, 2010). We previously noted the absence of hydrogenase sequences phylogenetically linked with these bacteria (Brazelton, Nelson & Schrenk, 2012), but recent sequencing of additional genomes from *Thiomicrospira, Thiomicrorhabdus*, and *Hydrogenovibrio* (Scott et al., 2018) has revealed that many of the hydrogenase sequences in Lost City metagenomes are affiliated with these taxa after all.

[FeFe]-hydrogenases typically associated with the production of H_2_ during fermentation were represented by HndCD sequences in several MAGs (**Figure 4**). This hydrogenase is capable of H_2_ oxidation with reduction of NADP in some organisms (Kpebe et al., 2018), but the presence of only one subunit in multiple Lost City MAGs (**Figure 4**) is curious and has unknown implications for the ability of these organisms to either consume or produce H_2_.

### Formate dehydrogenase and transporters

Formate dehydrogenase (FDH) catalyzes the reversible oxidation of formate to carbon dioxide, and various forms of FDH have diverse physiological roles in all three domains of life (Maia, Moura & Moura, 2015). Oxidation of formate was detected in all Lost City fluid samples, including those with significant contributions from ambient seawater (**Supplemental Table S8**).

We identified at least three kinds of FDH in Lost City fluids plus two distinct variants of FDH-like sequences. (1) NAD(P)-dependent FDH catalyzing formate oxidation in bacteria (K00123; FdoG/FdhF/FdwA) was detected in *Thermodesulfovibrionales* and WOR-3 MAGs. (2) NAD(P)-dependent FDH catalyzing reduction of carbon dioxide into formate (K05299; FdhA) was detected in Bipolaricaulota, *Thermodesulfovibrionales*, *Desulfotomaculum*, and *Dehalococcoidia* MAGs. (3) F_420_-dependent FDH catalyzing formate oxidation in methanogens (FdhA) was detected in the *Methanosarcinaceae* MAG. (4) A divergent FDH-like sequence was detected in *Methanosarcinaceae*, ANME-1, Bipolaricaulota, and *Thermodesulfovibrionales* MAGs. (5) Another divergent FDH-like sequence was detected in the WOR-3 MAG (**Figure 4**).

The divergent FDH-like sequence shared by the *Methanosarcinaceae*, ANME-1, Bipolaricaulota, and *Thermodesulfovibrionales* MAGs is also found in MAGs and SAGs (single-amplified genomes) from three continental serpentinite-hosted springs: the Voltri Massif in Italy (Brazelton et al., 2017), The Cedars in California, USA (Suzuki, Nealson & Ishii, 2018), and Hakuba Happo hot springs in Japan (Merino et al., 2020) (**Figure 6; Supplemental Figure S9**). All of these sequences from serpentinizing systems are more similar to each other than to any other sequences in public databases.

**Figure 6.**
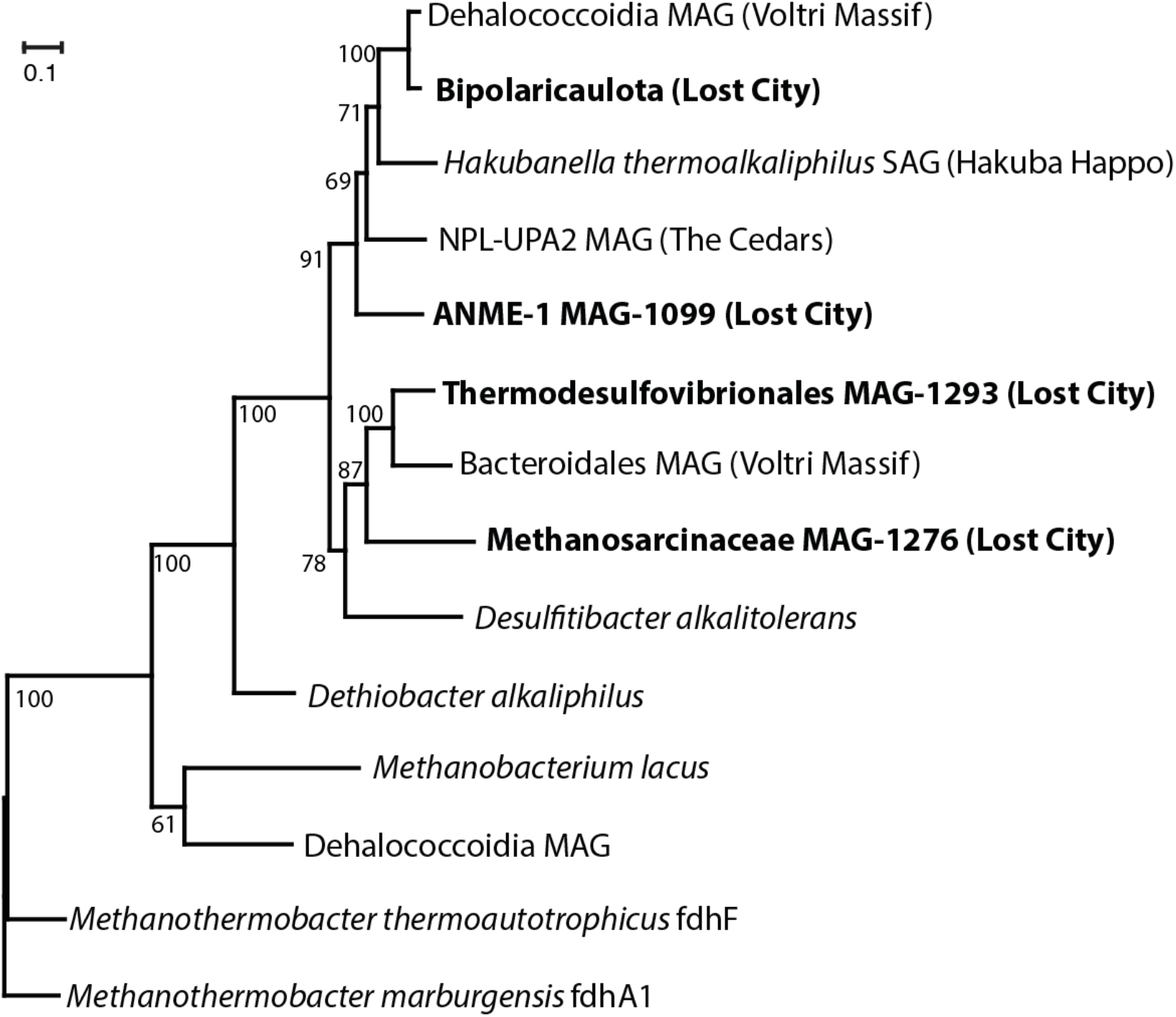
Phylogeny of divergent FDH-like sequences. Sequences identified in refined MAGs from this study are highlighted in bold font. Their closest matches in the NCBI nr database are from other serpentinite-hosted springs (Voltri Massif, Hakuba Happo, and The Cedars). The FDH-like sequences shown here include an iron-sulfur binding domain and a molybdopterin oxidoreductase domain, which are encoded as two separate coding regions in some species and as a fused coding region in others (see **Supplemental Figure S4 for** an expanded version of this figure including genomic context). The phylogeny was constructed from the conserved oxidoreductase domain. Bootstrap support values are shown for each node. The Lost City Bipolaricaulota sequence was identified in multiple BinSanity bins classified as Bipolaricaulota, but it was not included in the final, manually refined MAGs.

The formate transporters FdhC and FocA that were previously identified in Lost City chimney biofilms (Lang et al., 2018; McGonigle, Lang & Brazelton, 2020) were also detected in the metagenomes of venting fluids reported here, but they were only present at very low coverage (**Supplemental Table S7**), suggesting that they are specific to organisms inhabiting chimney biofilms. None of the MAGs highlighted by this study contain any known formate transporters. A lack of canonical formate transporters was also reported recently for a formate-utilizing methanogen in serpentinite-hosted, hyperalkaline groundwaters (Fones et al., 2021). Therefore, transport of formate into the cells of organisms inhabiting hyperalkaline subsurface environments may be carried out by uncharacterized proteins.

### Sulfate reduction

Surprisingly, the samples of sulfidic fluids collected from the chimney at Marker 2 (**Table 1**) did not contain elevated levels of taxa expected to represent sulfate-reducing bacteria (SRB) (**Figures 2-3**) or the genes encoding dissimilatory sulfite reductase (DsrAB) (**Figure 7**). Instead, Marker 2 fluids are dominated by aerobic bacteria that are likely to be adapted to chimney biofilms or to shallow subsurface zones with exposure to ambient seawater. Potential SRB such as *Thermodesulfovibrionales* were most abundant in the fluids venting from Marker 3, Marker C, Sombrero, and Calypso (**Figures 2-3**).

**Figure 7.**
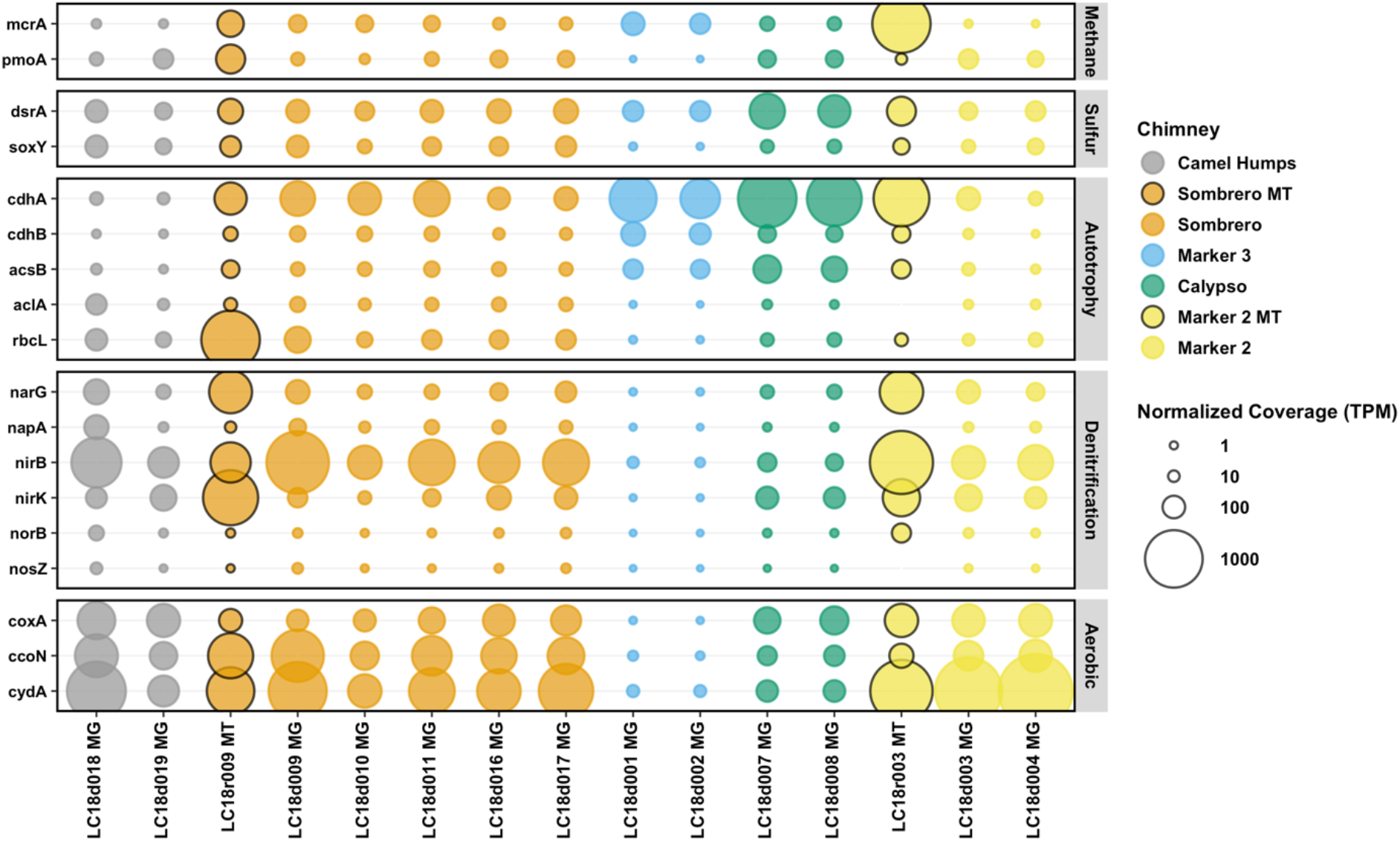
Abundance of key genes in Lost City hydrothermal fluid samples. Metagenomic coverage was normalized to predicted protein length and to the size of the metagenome or metatranscriptome library. The final normalized coverage is reported as a proportional unit (transcripts/fragments per million; TPM) suitable for cross-sample comparisons. Bubbles representing coverage in metatranscriptomes (MT), rather than metagenomes (MG), are highlighted with black borders. Genes are defined with KEGG Orthology; see **Supplemental Table S5**.

The other potential SRB in Lost City fluids include *Desulfotomaculum, Desulfocapsa,* and *Desulfobulbus. Desulfotomaculum* have been implicated as potential SRB in Lost City chimney biofilms (Gerasimchuk et al., 2010), but the *Desulfotomaculum* MAGs have neither hydrogenases nor carbon fixation enzymes, so their ability to reduce sulfate is dependent on the availability of organic matter. Furthermore, some *Desulfotomaculum* species are known to be incapable of sulfate reduction despite encoding DsrAB (Imachi et al., 2006). They do encode the nitrogenase enzyme required for nitrogen fixation, as do the *Methanosarcinaceae*, ANME-1, and *Thermodesulfovibrionales* MAGs (**Figure 4**). *Desulfobulbus* sequences were very rare in fluids from Markers 3 and C. *Desulfocapsa* were moderately abundant in Marker 3 fluids, but no MAGs classified as *Desulfocapsa* could be recovered during this study. Additionally, most of the dsrAB sequences in Lost City fluids were affiliated with *Thermodesulfovibrionales* or *Desulfotomaculum*; no dsrAB sequences belonging to *Desulfocapsa* or *Desulfobulbus* were identified in high-coverage contigs.

### Methane oxidation

Methane is present in Lost City fluids at a remarkably constant concentration of ∼1 mM, while concentrations of H_2_, sulfate, sulfide, and other chemicals vary widely (Kelley et al., 2005; Lang et al., 2012; Aquino et al., In Revision). The source of the methane, i.e. whether it is synthesized abiotically as a product of serpentinization or released from carbon stored within basement rocks, remains uncertain (Kelley & Früh-Green, 1999; Wang et al., 2018; Klein, Grozeva & Seewald, 2019; Labidi et al., 2020). Oxidation of methane was detected in most Lost City fluid samples, except the sample of Marker 3 fluids (**Supplemental Table S8**).

The primary candidates for the anaerobic oxidation of methane at Lost City are the ANME-1 archaea, which are most abundant in Calypso fluids (**Figures 2-3**). The absence of cytochromes and presence of hydrogenases in the ANME-1 MAG was noted by (Chadwick et al., 2021) as consistent with the genomic features of the so-called “freshwater” clade of ANME-1, for which the genus “Candidatus Methanoalium” was proposed. One of the shared features within this clade, including the Lost City ANME-1 MAG, is a novel HdrABC-MvhADG complex, which is involved in the transfer of electrons derived from H_2_ in methanogens. Therefore, this clade of ANME-1 may be involved in the H_2_-fueled production of methane instead of, or in addition to, the oxidation of methane. Distinguishing between methanogenesis and the anaerobic oxidation of methane with genomic data alone is notoriously difficult (Chadwick et al., 2021), and the *Methanosarcinaceae* and ANME-1 MAGs reported here contain features that are potentially consistent with both the production and oxidation of methane.

Potential methanotrophic bacteria were represented by ASVs classified as the gammaproteobacterial family *Methylomonaceae* (e.g. *Methylobacterium*), but they are expected to represent chimney biofilm communities (Brazelton et al., 2006) and were not abundant in any of the fluids included in this study. ASVs classified as *Methyloceanibacter*, various species of which can aerobically oxidize methane, methanol, or other methylated compounds (Vekeman et al., 2016), were prominent in Marker C fluids and very rare or absent in all other fluids (**Supplemental Table S2**).

### Carbonic anhydrase

At the high pH conditions of Lost City fluids, dissolved bicarbonate and carbonate are more stable than carbon dioxide, and the potential use of bicarbonate or carbonate as carbon sources has been explored in studies of continental sites of serpentinization (Suzuki et al., 2014, 2017; Kohl et al., 2016; Miller et al., 2018; Kraus et al., 2021; Fones et al., 2021). Carbonic anhydrase catalyzes the reversible conversion between bicarbonate and carbon dioxide, which may enable cells to utilize bicarbonate obtained from the environment. *Methanosarcinaceae* MAG-1276 encodes a carbonic anhydrase that shares 59-85% amino acid identities with sequences found in three other MAGs from Lost City (classified as NPL-UPA2 and Bipolaricaulota) and in one MAG from the Hakuba Happo hot spring (Nobu et al., 2021). These novel carbonic anhydrase sequences share only 35-41% amino acid identities with previously characterized proteins, e.g. the beta class carbonic anhydrases from *Clostridium aceticum* (**Supplemental Figure S10**). The Lost City carbonic anhydrase sequences retain each of the conserved residues highlighted by (Smith & Ferry, 2000) for beta class carbonic anhydrases. In addition, the *Methanosarcinaceae* MAG includes a predicted high-affinity bicarbonate transporter (SbtA).

### Glycine reductase

Glycine may be generated abiotically in high-H_2_ conditions or released as a primary thermogenic degradation production of biomass (Amend & Shock, 1998; Aubrey, Cleaves & Bada, 2009; Lang et al., 2013; Dick & Shock, 2021). The reduction of glycine to acetyl-phosphate is catalyzed by glycine reductase, which has been identified in metagenomes from multiple serpentinite-hosted springs (Nobu et al., 2021). Seven of the Lost City MAGs encode glycine reductase, and in most of these genomes, glycine reductase (GrdEBCA) is in a gene cluster that includes selenium transferase (SelA), selenocysteine-specific elongation factor (SelB), and thioredoxin (TrxA) (**Supplemental Table S5**), consistent with the gene organization of bacteria that conserve energy by reduction of glycine (Andreesen, 2004). Each of these MAGs also encodes partial Wood-Ljungdahl pathways, suggesting that they may use glycine reductase as part of the reductive glycine pathway for carbon fixation (Sánchez-Andrea et al., 2020).

### ATP synthase

The production of ATP is catalyzed by the enzyme ATP synthase, which diverged into distinct archaeal and bacterial versions early in the evolution of life (Müller & Grüber, 2003). A few of the bacterial MAGs in this study encode the archaeal form of ATP synthase (A-type) instead of the bacterial form (F-type). These include *Dehalococcoidia* MAG-844, Paceibacteria MAG-855, WOR-3 MAG-1066, and all three NPL-UPA2 MAGs (**Supplemental Table S5**). Chloroflexi, Paceibacteria (previously named candidate phylum OD1), and NPL-UPA2 bacteria have also been observed to encode A-type ATP synthase in The Cedars, a continental serpentinite spring (Suzuki et al., 2017; Suzuki, Nealson & Ishii, 2018), suggesting that the A-type ATP synthase may provide advantages to both bacteria and archaea in the high pH, highly reducing conditions created by serpentinization.

In addition, ATP synthase was completely absent in three of the Paceibacteria MAGs, as was the case for multiple Paceibacteria MAGs from The Cedars (Suzuki et al., 2017). *Natronincolaceae* MAG-1138 also lacks any ATP synthase genes, and its genomic content suggests an obligate fermentative lifestyle (**Supplemental Material**). Other genera within family *Natronincolaceae* include *Alkaliphilus* and *Serpentinicella*, which have been isolated from the Prony Bay hydrothermal field (Mei et al., 2016; Postec et al., 2021).

## DISCUSSION

### Distinct zones of microbial activity in Lost City’s basement

The massive edifice of Poseidon towers 60 meters above the center of the Lost City hydrothermal field (**Figure 1**). Alkaline hydrothermal fluids flow from the serpentinite basement and throughout the Poseidon structure, exiting at multiple locations across the field. The differing flow paths that lead to each location have distinct residence times (Moore et al., 2021) and produce distinct chemical and microbiological compositions of the venting fluids (Lang et al., 2012, 2018, 2021; Aquino et al., In Revision).

For example, the venting locations Marker 3 and Camel Humps sit only a few meters from each other at the summit of Poseidon, but the fluids venting from each structure appear to have taken different paths, which is reflected in their distinct microbial communities. Marker 3 fluids are dominated by a few archaeal and bacterial species that have the genomic potential to metabolize H_2_, formate, and sulfate. Genes encoding methanogenesis, sulfate reduction, and carbon fixation are much more abundant in Marker 3 fluids than genes encoding aerobic respiration (**Figure 7; Supplemental Figures S12-S15**). In contrast, Camel Humps fluids host a diverse assemblage of bacteria capable of using oxygen, nitrate, and nitrite as oxidants. These taxonomic and metabolic patterns are generally similar between ribosomal gene and ribosomal RNA datasets (**Figure 2**) and between metagenomes and metatranscriptomes (**Figures 3 and 7**) from the same locations, indicating that the most abundant organisms in these fluids were likely to have been metabolically active at the time of sampling.

### Sulfate reduction is limited to a few taxa in the subseafloor

Previous studies of Lost City hydrothermal fluids have revealed a consistent trend across the field in which the consumption of H_2_ and sulfate is correlated with the production of hydrogen sulfide (Proskurowski et al., 2008; Lang et al., 2012, 2018). Therefore, sulfate-reducing bacteria (SRB) are expected to be widespread and metabolically active in the subsurface environments below the Lost City chimneys.

The metagenomic results presented here indicate a single, novel species of *Thermodesulfovibrionales* as the SRB that is most likely to be responsible for these trends. It dominates the fluids at Marker C, Marker 3, and Calypso, and it accounts for most of the genes associated with sulfate reduction and H_2_ oxidation in these fluids. It also includes multiple formate dehydrogenases and various genes indicative of organic carbon oxidation (**Supplemental Table S5**), suggesting metabolic flexibility that is not dependent on the availability of H_2_ and inorganic carbon.

The Lost City *Thermodesulfovibrionales* belong to a novel clade associated with deep subsurface environments and hot springs that shares only 82-87% nucleotide identities with characterized *Thermodesulfovibrio* species (**Supplemental Figure S7**). This clade also includes a 16S rRNA sequence from highly alkaline borehole fluids associated with serpentinization of the Samail Ophiolite in Oman (Rempfert et al., 2017). *Thermodesulfovibrionales* are not abundant in other sites of serpentinization, although DsrB sequences with similarity to *Thermodesulfovibrio* were detected in alkaline borehole fluids from the Coast Range Ophiolite (Sabuda et al., 2020). Sulfate concentrations are much higher in borehole fluids from the Samail Ophiolite (up to 3.9 mM) and the Coast Range Ophiolite (up to 0.4 mM) compared to most natural springs associated with serpentinization (e.g. <0.02 mM in the Tablelands, Voltri Massif, and The Cedars) (Brazelton et al., 2017; Rempfert et al., 2017; Sabuda et al., 2020; Cook et al., 2021). An exception is Ney Springs, where sulfate can be as high as 12.9 mM, but the potential SRB detected there did not include *Thermodesulfovibrionales* (Trutschel et al., In Revision).

### H*_2_*-fueled metabolism is limited to a few taxa in the subseafloor

Lost City fluids contain copious quantities of H_2_ (1-7 mM, with subsurface concentrations predicted to reach 14 mM; (Kelley et al., 2005; Aquino et al., In Revision), which is expected to be a tremendous boost to life in the subseafloor. Surprisingly, only two taxa (*Thermodesulfovibrionales* and *Methanosarcinaceae)* that are abundant in Lost City fluids encode hydrogenases known to be associated with H_2_ oxidation. Therefore, the ability of the subseafloor ecosystem to be powered by H_2_ may depend on one species of bacteria and one species of archaea.

Another type of hydrogenase, known as membrane-bound hydrogenase (Mbh), was also detected in *Methanosarcinaceae*, ANME-1, and Bipolaricaulota genomes (**Figure 5**). In *Thermococcus* and *Pyrococcus*, Mbh is responsible for H_2_ production during anaerobic, heterotrophic growth, and some bacteria use Mbh in coordination with FDH to convert formate into H_2_ (Schut et al., 2013; Nobu et al., 2015). In heterotrophic Bipolaricaulota, Mbh has been proposed to couple the production of H_2_ with ATP synthesis in coordination with the MvhAGD-HdrABC complex (Youssef et al., 2019). In methanogens, the role of Mbh is unclear, but each of the methanogens that encode Mbh can utilize either formate or H_2_ as their sole source of electrons. In H_2_-saturated Lost City fluids, biological production of additional H_2_ seems highly unfavorable, and the sequence divergence between the Lost City sequences and these previously characterized Mbh prevents any firm conclusions on whether they are more likely to catalyze the consumption or production of H_2_.

### Formate metabolism may operate via unknown mechanisms in the subseafloor

Formate forms abiotically in the high-pH, reducing conditions of serpentinizing fluids, and it is the second-most abundant form of carbon in Lost City fluids after methane and the second-most available reductant after dissolved H_2_ (Lang & Brazelton, 2020). Much of the biomass in Lost City chimneys is produced from formate that is derived from carbon originating in Earth’s mantle (Lang et al., 2018). Formate is the preferred substrate for methanogens in at least one other site of serpentinization where carbon dioxide is limiting (Fones et al., 2021). However, none of the taxa highlighted by this study contain any known formate transporters, and surprisingly few encode formate dehydrogenase (FDH), the enzyme that catalyzes the oxidation of formate. A remarkable exception is *Thermodesulfovibrionales*, which encodes three distinct forms of FDH.

A divergent, FDH-like sequence with unknown function was shared by four of the key taxa in this study (*Thermodesulfovibrionales, Methanosarcinacae,* ANME-1, and Bipolaricaulota). These sequences form a distinct clade that includes sequences from continental serpentinite springs, suggesting that this gene represents a shared, unknown metabolic strategy in serpentinizing fluids (**Figure 6**).

In the highly reducing conditions of Lost City fluids, biosynthetic pathways are more energetically favorable than in typical environments, and the synthesis of some biomolecules can even be energy-yielding (Amend et al., 2011; Dick & Shock, 2021). Therefore, the ability to incorporate formate directly into metabolic pathways, rather than first oxidizing it to carbon dioxide, could be a competitive advantage in Lost City’s basement, where formate is 100-1,000 times more abundant than carbon dioxide (Lang & Brazelton, 2020). Potential evidence for this hypothesis is the prevalence of partial and complete Wood-Ljungdahl pathways among Lost City bacteria (**Supplemental Table S5**). Eight of these genomes do not encode a known FDH, suggesting that they may be able to use formate, rather than carbon dioxide, as the substrate for carbon fixation and perhaps acetogenesis. Some acetogens can use formate as their sole source of energy and carbon, although FDH may be still required to supply carbon dioxide as an electron acceptor (Jain et al., 2020).

In the absence of FDH, pyruvate formate lyase (PflD), which is encoded by some of the same genomes with partial Wood-Ljungdahl pathways (Bipolaricaulota, NPL-UPA2, and *Dehalococcoidia*), might catalyze the reduction of formate directly into acetyl-CoA and pyruvate (Zelcbuch et al., 2016). However, this activity has only been demonstrated in *E. coli*, and its relevance to these taxa in the unusual environmental conditions of Lost City requires further research.

## Conclusions

This study has highlighted multiple examples of metabolic strategies shared among the archaea and bacteria most likely to inhabit subsurface habitats underlying the Lost City hydrothermal field. These shared strategies appear to be advantageous for life in environments that are rich in H_2_ (e.g., hydrogenases), provide a steady supply of simple organic molecules (e.g., formate dehydrogenase, pyruvate formate lyase, and glycine reductase), lack carbon dioxide (e.g., carbonic anhydrase), and make typical ATP synthesis too difficult or unnecessary.

Many of the predicted proteins associated with these metabolic strategies are not closely related to any previously characterized enzymes, but they are shared by diverse archaea and bacteria in Lost City and other sites of serpentinization (e.g., Prony Bay, The Cedars, Hakuba Happo, and Voltri Massif), strongly suggesting the influence of horizontal gene transfer among these systems. The functions of these proteins are mostly unknown and require further study, but the results presented here indicate that they are likely to be important clues for understanding the ecology, physiology, and evolution of microbes adapted to these conditions.

If potential extraterrestrial habitats are evaluated for their ability to support a robust ecosystem over geological time scales (Cabrol, 2018), then it is critical to identify and understand the metabolic pathways of key organisms that form the foundations of ecosystems that are potentially relevant for astrobiology. All ecosystems on the surface of the Earth are based on autotrophs that rely on the availability of sunlight and carbon dioxide. The most promising extraterrestrial habitats in our solar system (Schulte et al., 2006; Waite et al., 2017; Jones, Goordial & Orcutt, 2018; Michalski et al., 2018), however, are dark, rock-hosted environments where simple organic molecules may be more biologically available than carbon dioxide. The organisms and metabolic pathways highlighted by this study can help us to understand the biological advantages and limitations of such conditions.

## METHODS

### Collection of hydrothermal fluid samples

Hydrothermal fluid samples were collected from actively venting chimneys at the Lost City hydrothermal field (**Figure 1**) using ROV *Jason* during the 2018 Lost City expedition aboard R/V *Atlantis* (AT42-01). On the seafloor, venting fluids were slowly pumped through 0.2 μm Millipore Sterivex cartridge filters or into acid-washed Kynar bags with the HOG sampler (Lang & Benitez-Nelson, 2021). Samples intended for RNA extractions were collected into 2 L Kynar bags containing 67 mL of a stop solution (97.5% 200 proof ethanol, 2.5% Trizol LS; Thermo Fisher). Fluid temperatures were monitored in real-time during sampling with a probe embedded into the sampler intake. Concentrations of sulfate, hydrogen sulfide, and magnesium were measured according to standard methods (Butterfield & Massoth, 1994). Additional sampling methods are provided in the **Supplemental Material**.

### Sequencing of DNA and RNA

Extraction of DNA and RNA from Sterivex filters was conducted as described previously (Brazelton et al., 2017; Thornton et al., 2020), with minor modifications described in the **Supplemental Material**. Sequencing of amplicons generated from 16S rRNA genes and cDNA was performed at the Genomics Core Facility at Michigan State University on an Illumina MiSeq instrument using dual-indexed Illumina fusion primers targeting the V4 region of the 16S rRNA gene (Kozich et al., 2013). Amplicon sequence variants (ASVs) were inferred from 16S rRNA amplicon sequences with DADA2 v. 1.10.1 (Callahan et al., 2016). Paired-end sequencing (2 x 125 bp) of metagenomic libraries was conducted at the University of Utah High-Throughput Genomics Core Facility at the Huntsman Cancer Institute with an Illumina HiSeq2500 platform. Metatranscriptome sequencing was conducted with a 150 cycle paired-end run on a NovaSeq 6000. Methods for the assembly and binning of metagenomes, including all downstream analyses are provided in the **Supplemental Material.**

### Data Availability

Amplicon sequences are available via NCBI BioProject PRJNA672129, and metagenome and metatranscriptome sequences are available via BioProject PRJNA779602. MAGs are associated with the same BioProject and are individually accessible via BioSamples SAMN23474158 - SAMN23474187. In addition, GenBank accessions are listed for each MAG in **Supplemental Table S4**. Protocols, metadata, and additional data are provided in a Zenodo-archived GitHub repository accessible via DOI: 10.5281/zenodo.5798015, and on the BCO-DMO page for project 658604: https://www.bco-dmo.org/award/658604.

## Acknowledgements

We thank the Scientific Party of the 2018 Lost City expedition (AT42-01), including the crews of R/V *Atlantis* and ROV *Jason*. In addition, we gratefully acknowledge invaluable mentorship for many years from John Baross and Deborah Kelley. Mitch Elend, Christopher Thornton, and Alex Hyer provided critical technical support before, during, and after the expedition. University of Utah students enrolled in BIOL 3270 / 5270 assisted in the analysis of metagenomic data. This work was supported by NSF awards to Brazelton and Lang (OCE-1536702/1536405), the NASA Astrobiology Institute Rock-Powered Life team, a NASA Postdoctoral fellowship to McGonigle, the Swiss National Science Foundation, and the Deep Carbon Observatory.

## Tables

1. Sample overview including temp and chemistry

## Figures

1. Map and ordination

2. 16S bubbles

3. MAG bubbles

4. MAG presence/absence of key genes

5. MBH phylogeny

6. FDH phylogeny

7. Key gene bubbles

## Supplemental Figures

1. Extended map figure

2. Chimney photos

3. Metagenome analysis workflow

4. Kaiju bubbles

5. Phylogeny – Methanosarcinales 16S + mcrA

6. Phylogeny – Bipolaricaulota 16S

7. Phylogeny – Thermodesulfovibrionales 16S

8. Phylogeny + gene order mbhL

9. Phylogeny + gene order FDH

10. Phylogeny – carbonic anhydrase

11. Phylogeny – GrdB

12. Hydrogenase bubbles

13. Acetate/formate bubbles

14. Methanogenesis bubbles

15. Acetogenesis bubbles

## Supplemental Tables (Excel files)

1. Sample info

2. Full 16S count table including contaminants

3. Kaiju tables

4. MAG taxonomy, completeness, coverage table

5. MAG gene presence absence tables

6. MAG annotations for transporters, dbCAN, FeGenie

7. KO coverage table

8. Incubation experiment results

## Github Repo

1. Protocols

2. Kaiju Krona plots

3. NCBI SRA and GenBank metadata

4. MAG sequences and annotations

5. Alignments and sequences for phylogenetic trees

6. R code for plots

7. Python scripts for metagenomic analyses

---

**Supplemental Figure S1.**
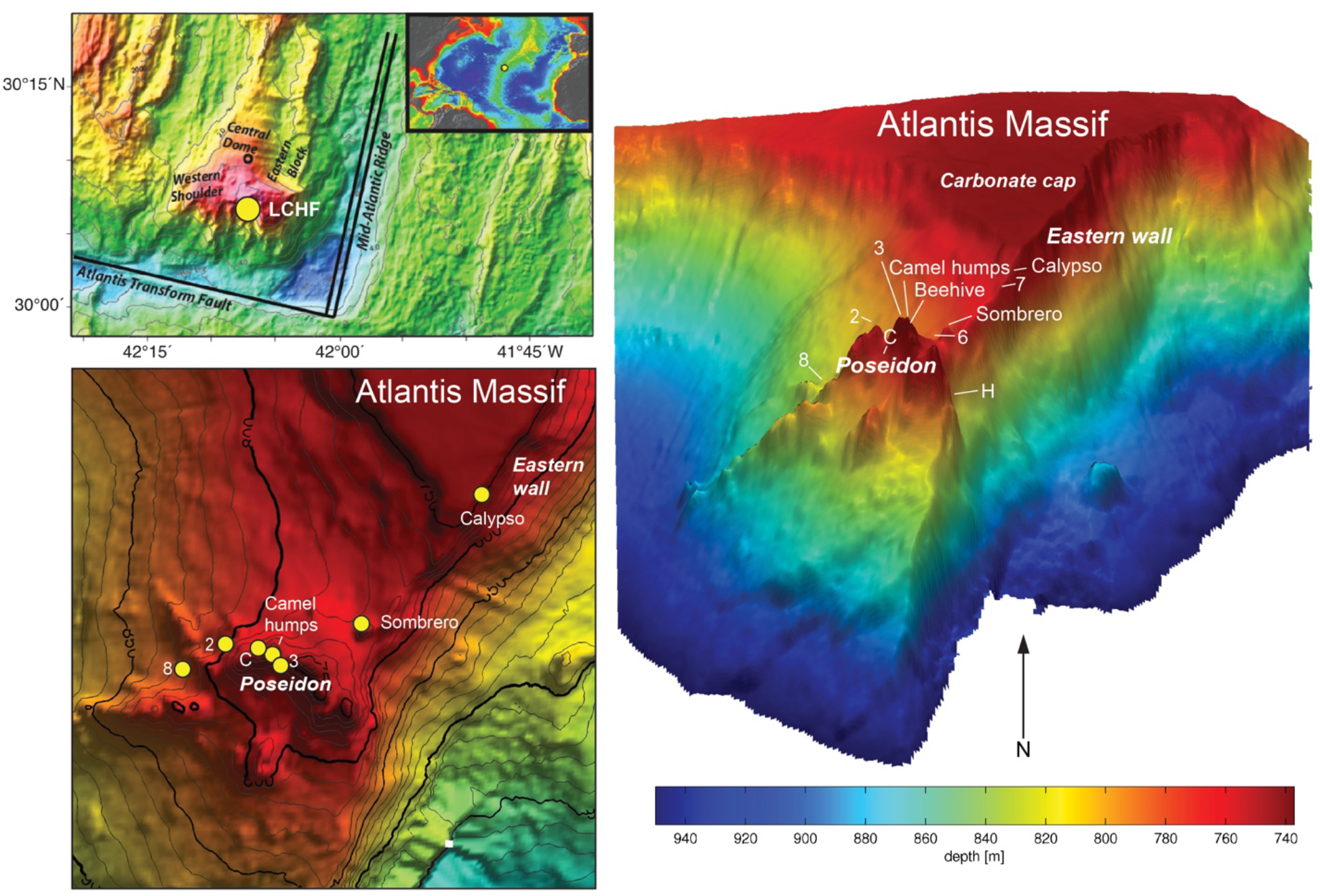
Extended version of **Figure 1** showing the location of the Lost City hydrothermal field near the summit of the Atlantis Massif, which is located northwest of the intersection of the Mid-Atlantic Ridge and the Atlantis Transform Fault.

**Supplemental Figure S2.**
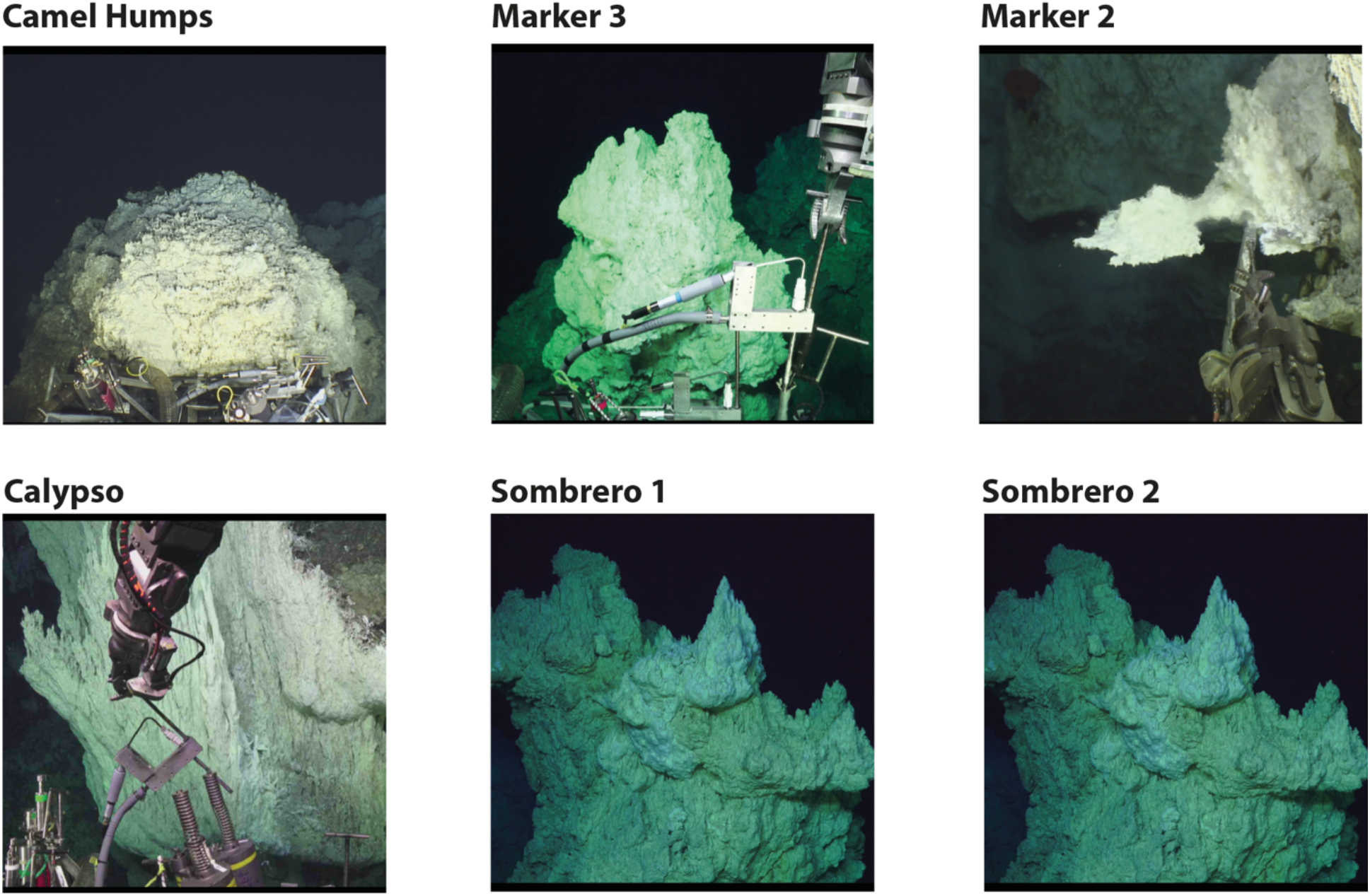
Photographs of sampling locations for this study, captured on the seafloor by ROV *Jason*. Camel Humps and Marker 3 are visually distinct structures despite their nearby locations. Sombrero was sampled at the same location on two separate dives.

**Supplemental Figure S3.**
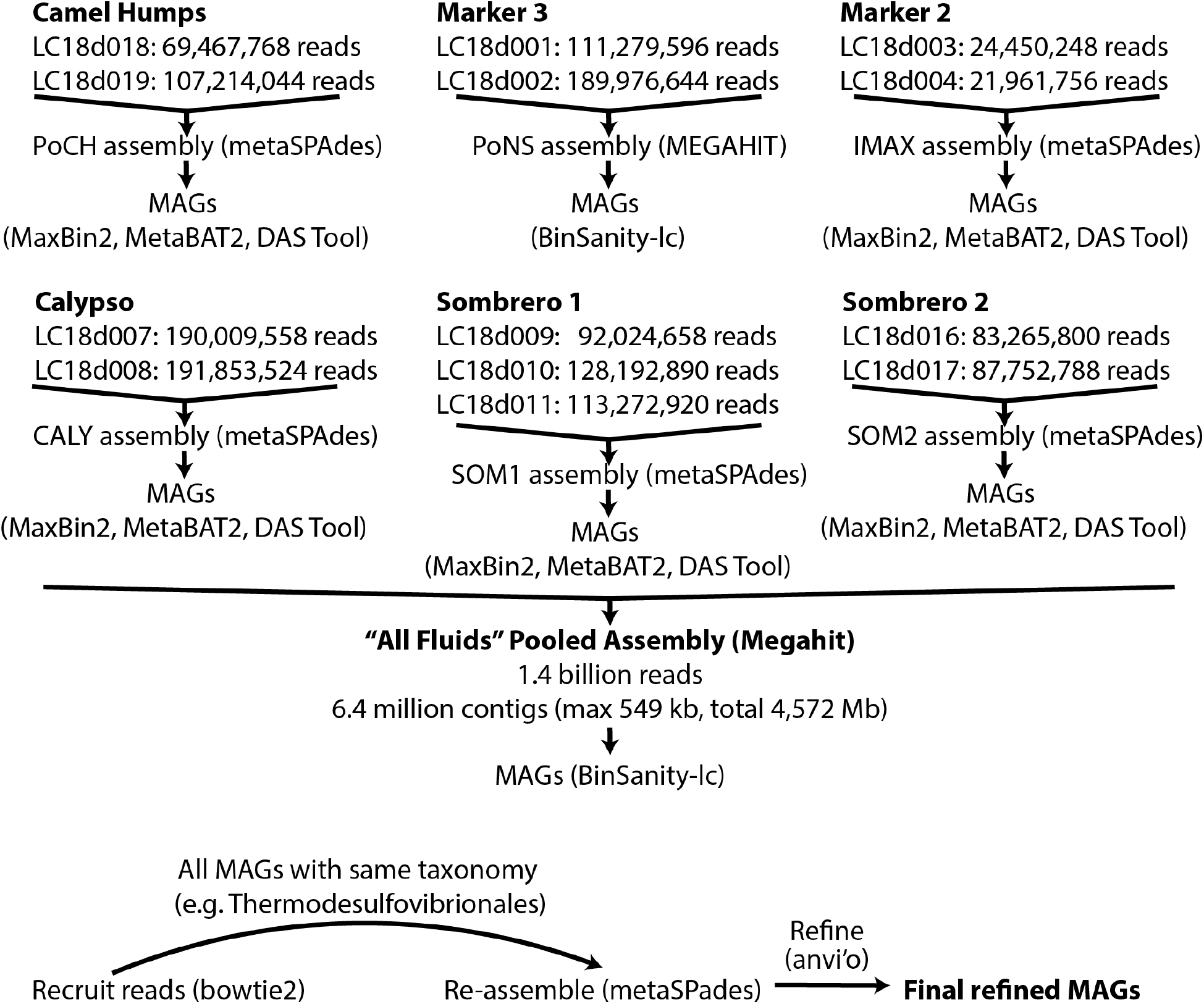
Overall workflow for assembly of metagenomes and binning into metagenome-assembled genomes (MAGs). Assemblies were performed with reads pooled from each chimney location (chimney-specific assemblies), and one “all fluids” pooled assembly was performed with all metagenomic reads from all chimney locations. Initial bins constructed with automated tools were used as a template for recruiting metagenomic reads for a re-assembly and manual curation and refinement of the final MAGs.

**Supplemental Figure S4.**
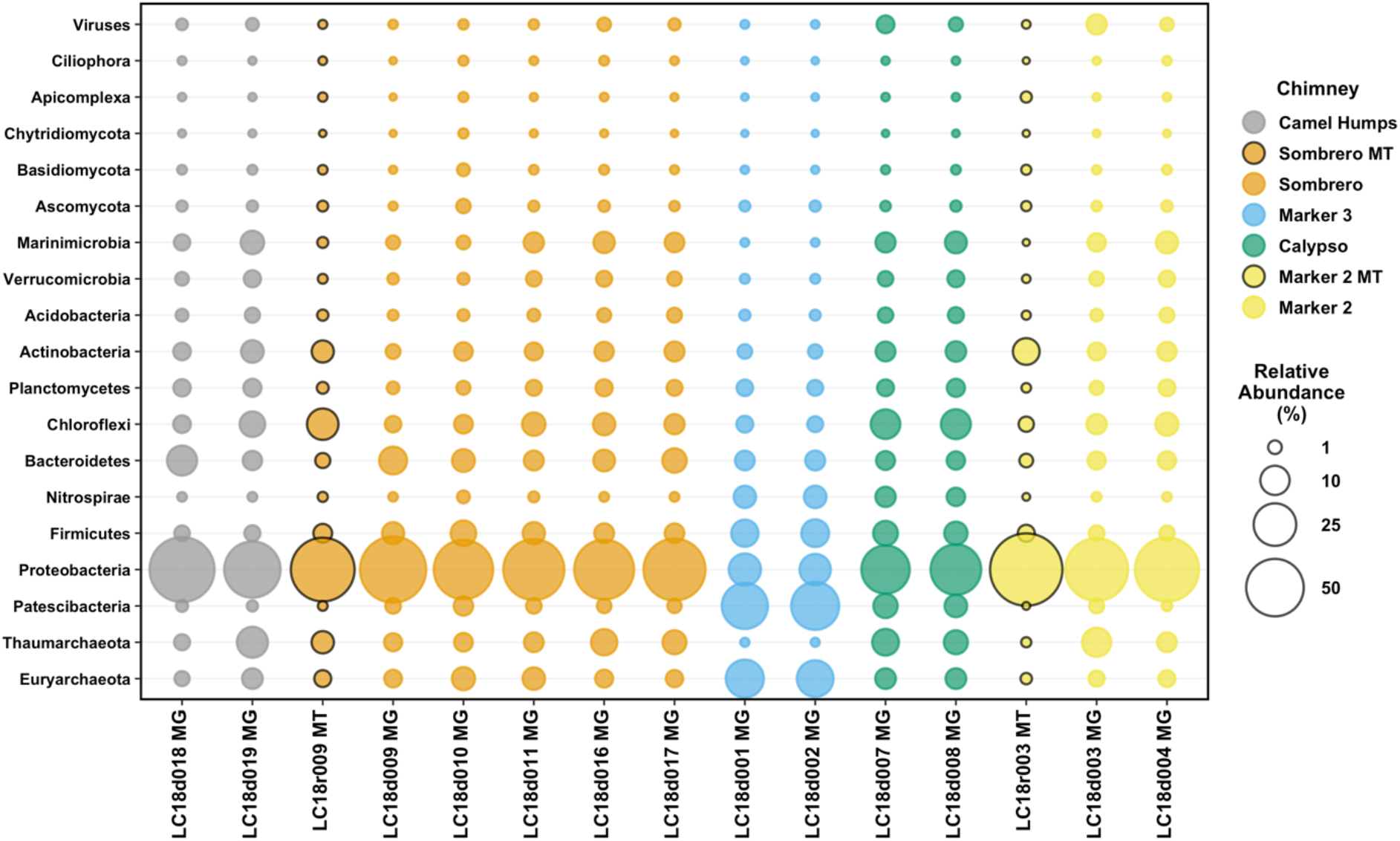
Percent of reads classified to the top 18 phyla (plus viruses) in Lost City hydrothermal fluid samples. Unassembled reads were classified using Kaiju with its default NCBI nr+euk database. Percentages were calculated as the number of reads classified to each phyla divided by the total number of reads in that library that could be classified to the phylum level by Kaiju. Bubbles representing reads in metatranscriptomes (MT), rather than metagenomes (MG), are highlighted with black borders.

**Supplemental Figure S5.**
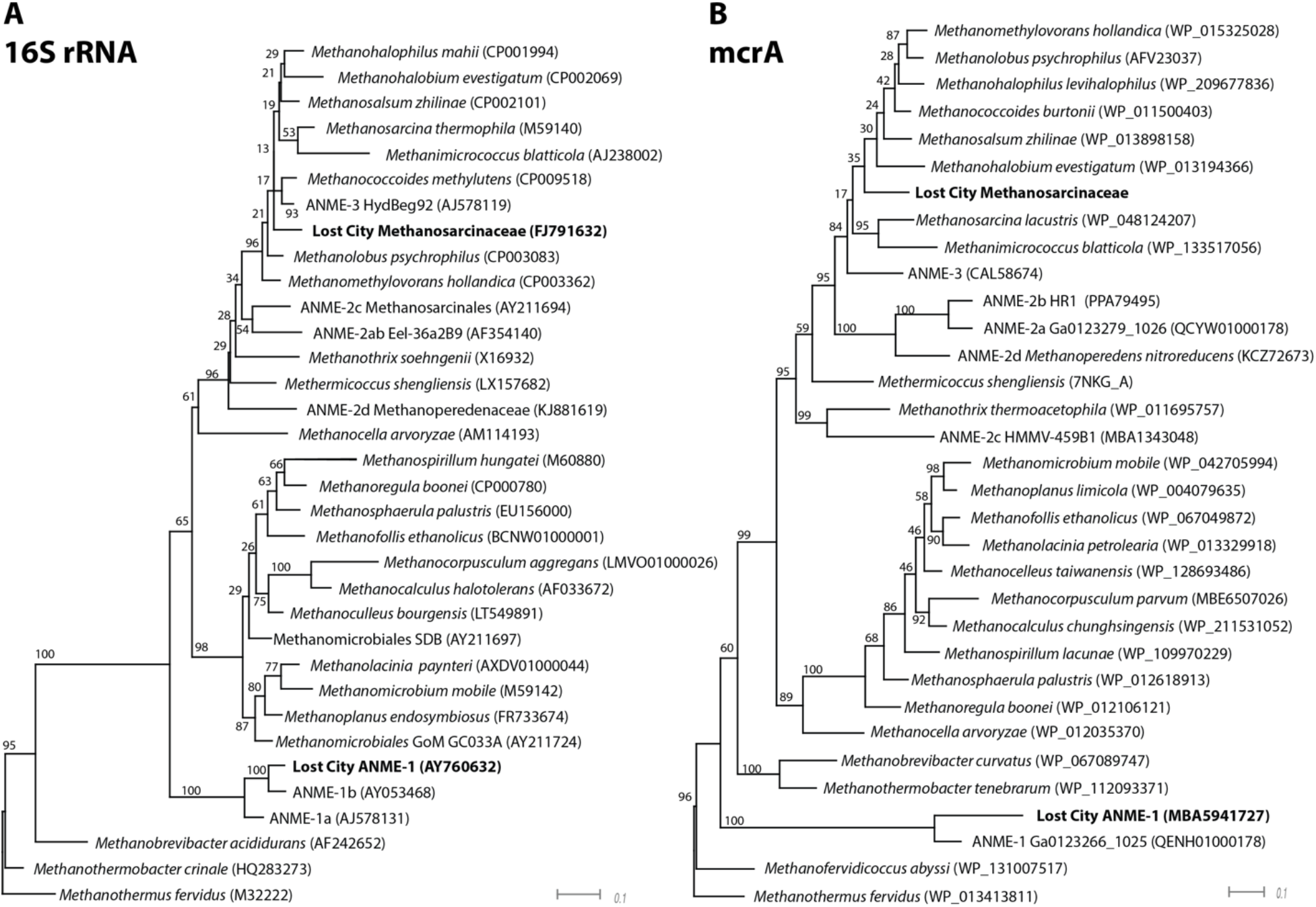
Phylogenies of 16S rRNA and mcrA (alpha subunit of methyl coenzyme M reductase) highlighting Lost City MAGs classified as Methanosarcinaceae and ANME-1. Bootstrap support values are shown for each node. Sequences and accession IDs are provided in the Zenodo-archived GitHub repository accessible via DOI: 10.5281/zenodo.5798015.

**Supplemental Figure S6.**
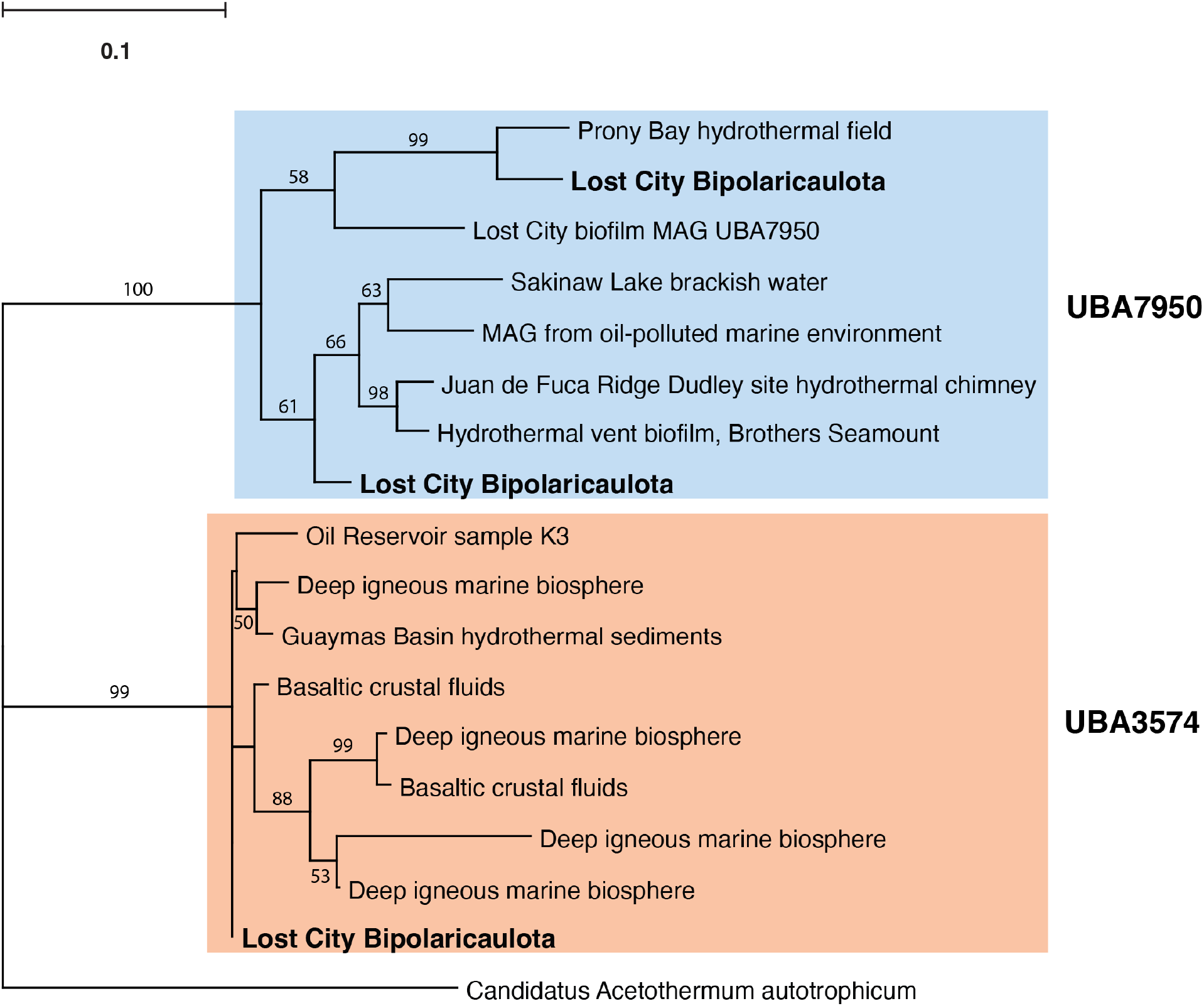
Phylogeny of 16S rRNA highlighting Lost City sequences classified as Bipolaricaulota. The most abundant Lost City Bipolaricaulota 16S rRNA sequences cluster into two distinct monophyletic groups, classified by GTDB as UBA3574 and UBA7950, which corresponds to the classifications of the three refined Bipolaricaulota MAGs (**Figure 3**). The UBA7950 sequences are further divided into two clades, one of which includes a MAG assembled by Parks et al. (2018) from our previous study of Lost City chimney biofilms (DOHL01000117). Bootstrap support values greater than 50 are shown for each node. Sequences and accession IDs are provided in the Zenodo-archived GitHub repository accessible via DOI: 10.5281/zenodo.5798015.

**Supplemental Figure S7.**
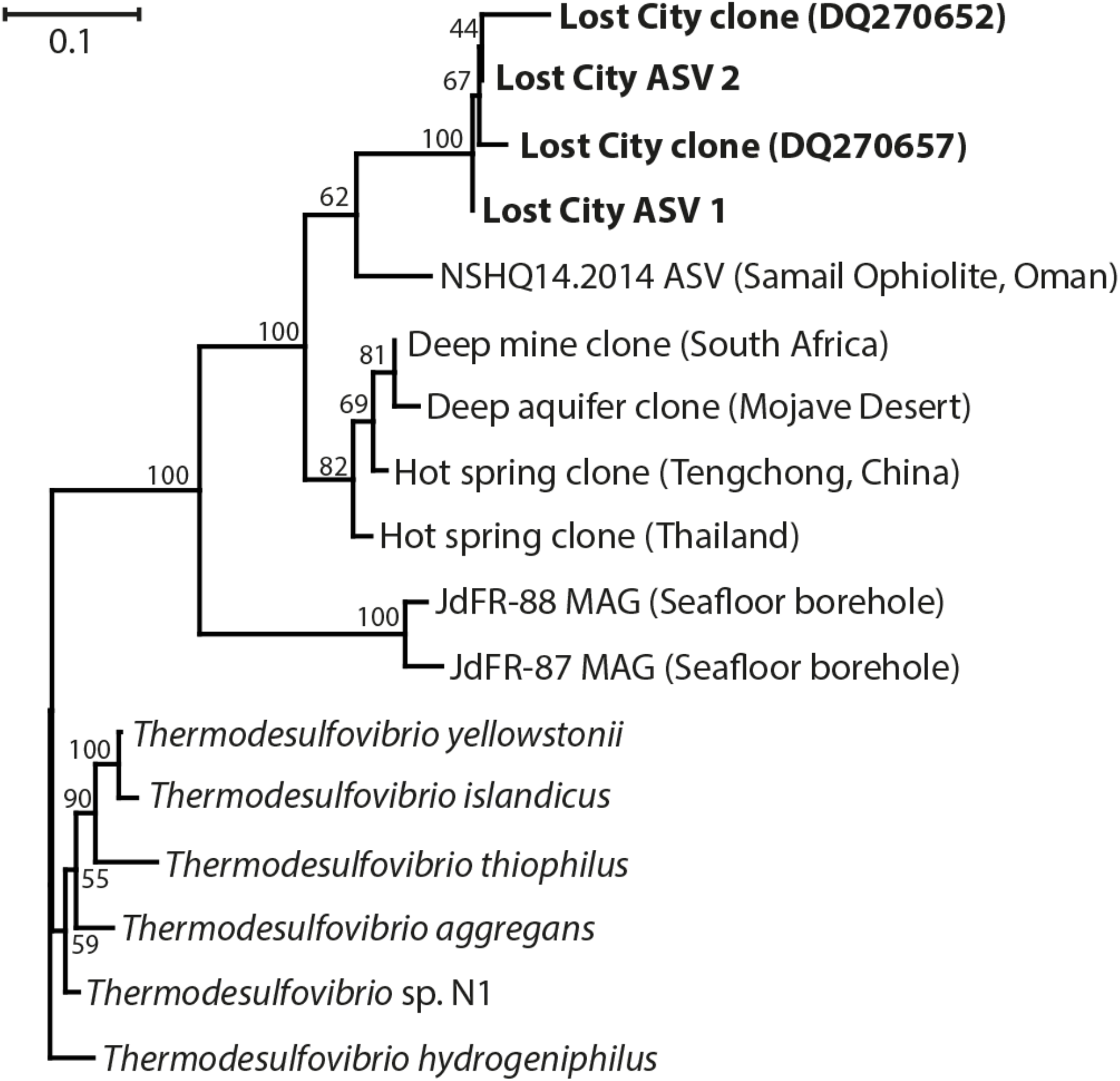
Phylogeny of 16S rRNA highlighting Lost City sequences classified as Thermodesulfovibrionia. The two Lost City ASVs differ from each other by a single base and match sequences from a previously published clone library of Lost City chimney biofilms (Brazelton et al., 2006). They share 90% nucleotide identities with their closest neighbor, an ASV from alkaline borehole fluids in the Samail Ophiolite, Oman (Rempfert et al., 2017). Sequences and accession IDs are provided in the Zenodo-archived GitHub repository accessible via DOI: 10.5281/zenodo.5798015.

**Supplemental Figure S8.**
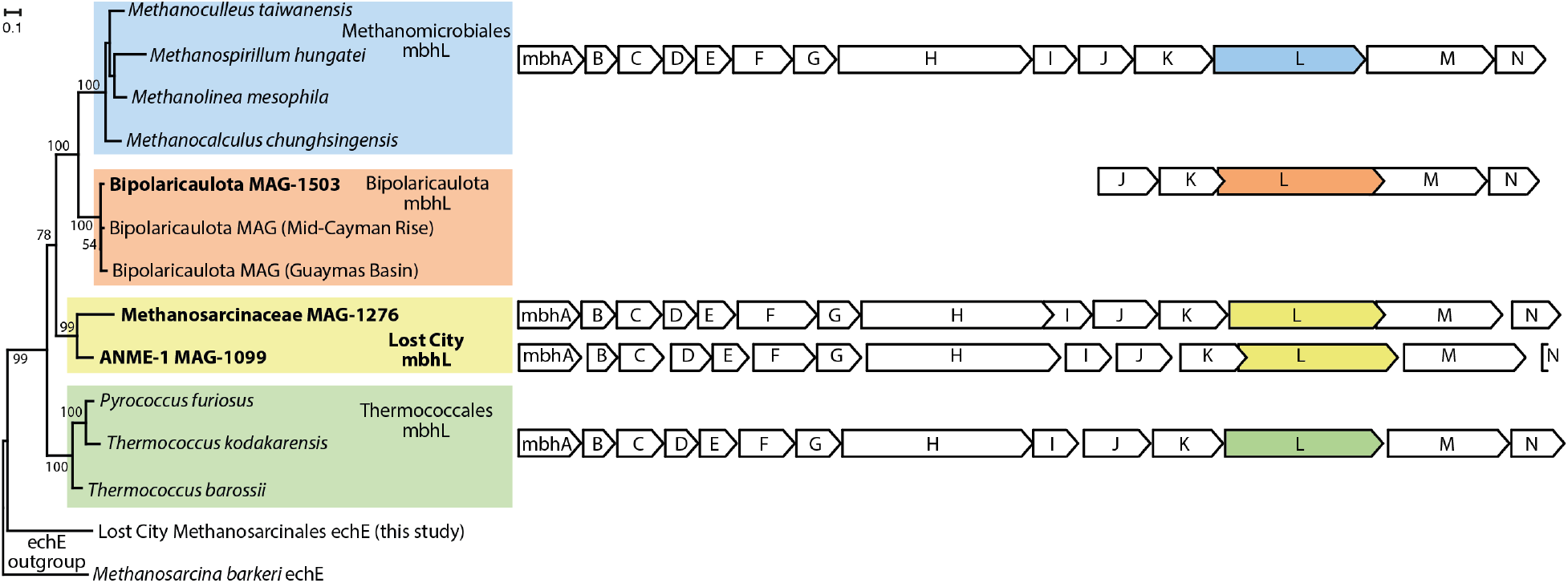
Phylogeny of the large catalytic subunit of membrane-bound hydrogenase (mbhL) and the mbh gene cluster (expanded version of Figure 5). Sequences and accession IDs are provided in the Zenodo-archived GitHub repository accessible via DOI: 10.5281/zenodo.5798015.

**Supplemental Figure S9.**
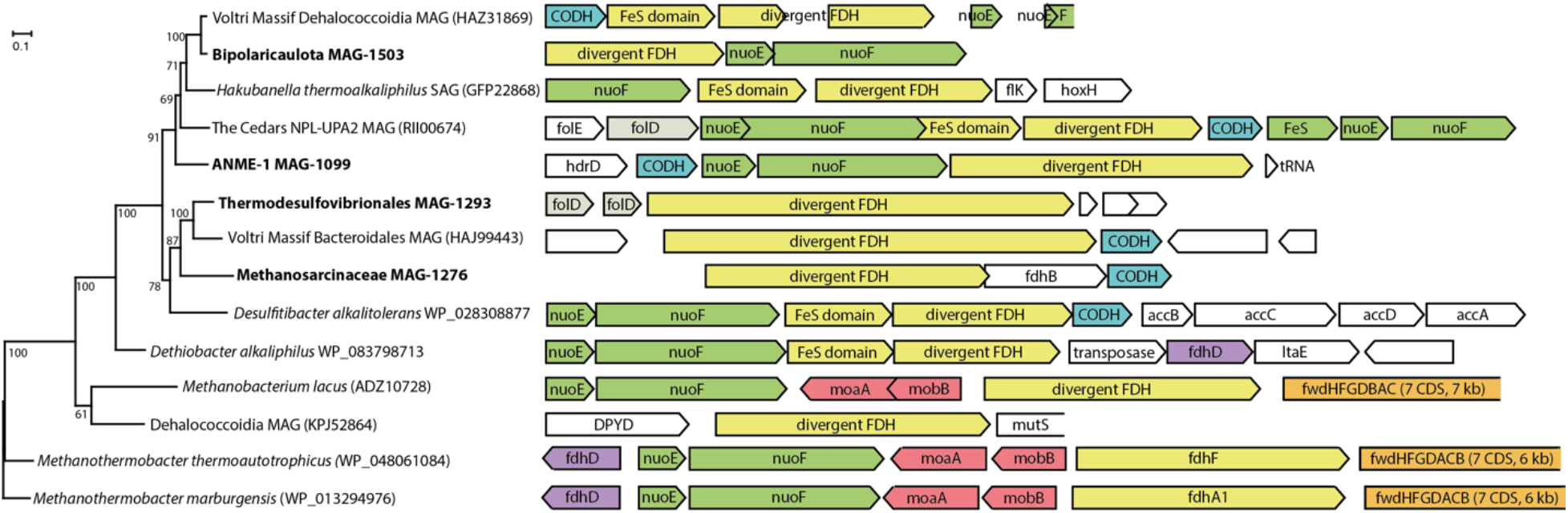
Phylogeny of divergent FDH-like sequences (expanded version of Figure 6). In most cases, the divergent FDH-like gene was flanked by nuoEF (encoding NADH-quinone oxidoreductase) and a hypothetical sequence with a conserved domain associated with monomeric carbon monoxide dehydrogenase (CODH). Furthermore, most of these gene clusters contained signs of genome instability just upstream or downstream such as pseudogenes, transposases, or a toxin/antitoxin system (not shown here). Sequences and accession IDs are provided in the Zenodo-archived GitHub repository accessible via DOI: 10.5281/zenodo.5798015.

**Supplemental Figure S10.**
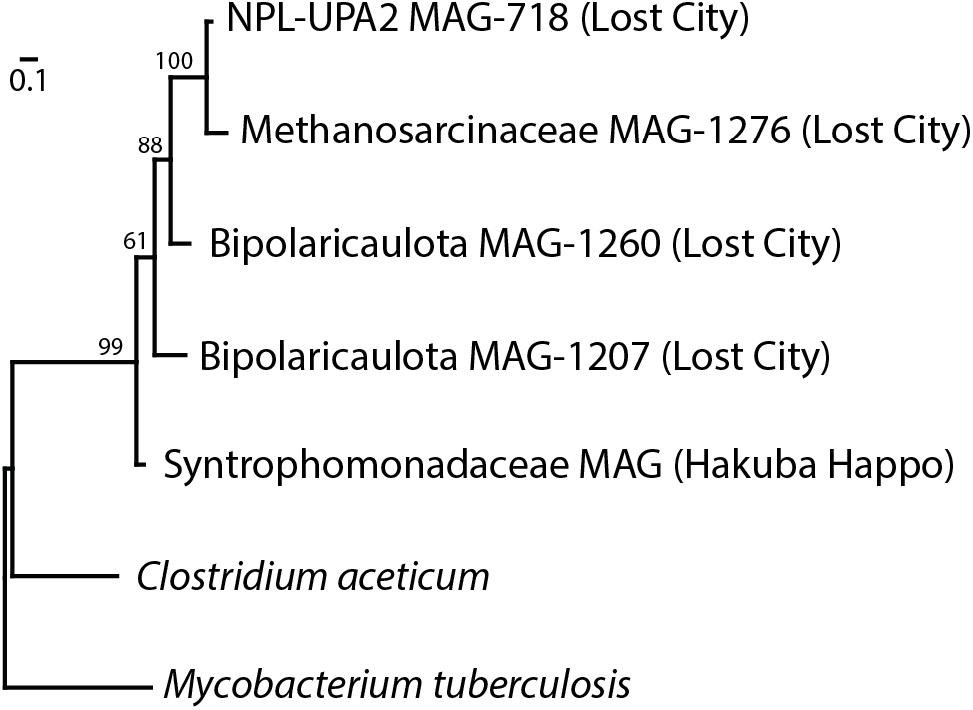
Phylogeny of divergent sequences predicted to encode carbonic anhydrase. Lost City sequences form a novel clade including a predicted sequence from a MAG recovered from another serpentinite-hosted spring (Hakuba Happo). Bootstrap support values are shown for each node. Sequences and accession IDs are provided in the Zenodo-archived GitHub repository accessible via DOI: 10.5281/zenodo.5798015.

**Supplemental Figure S11.**
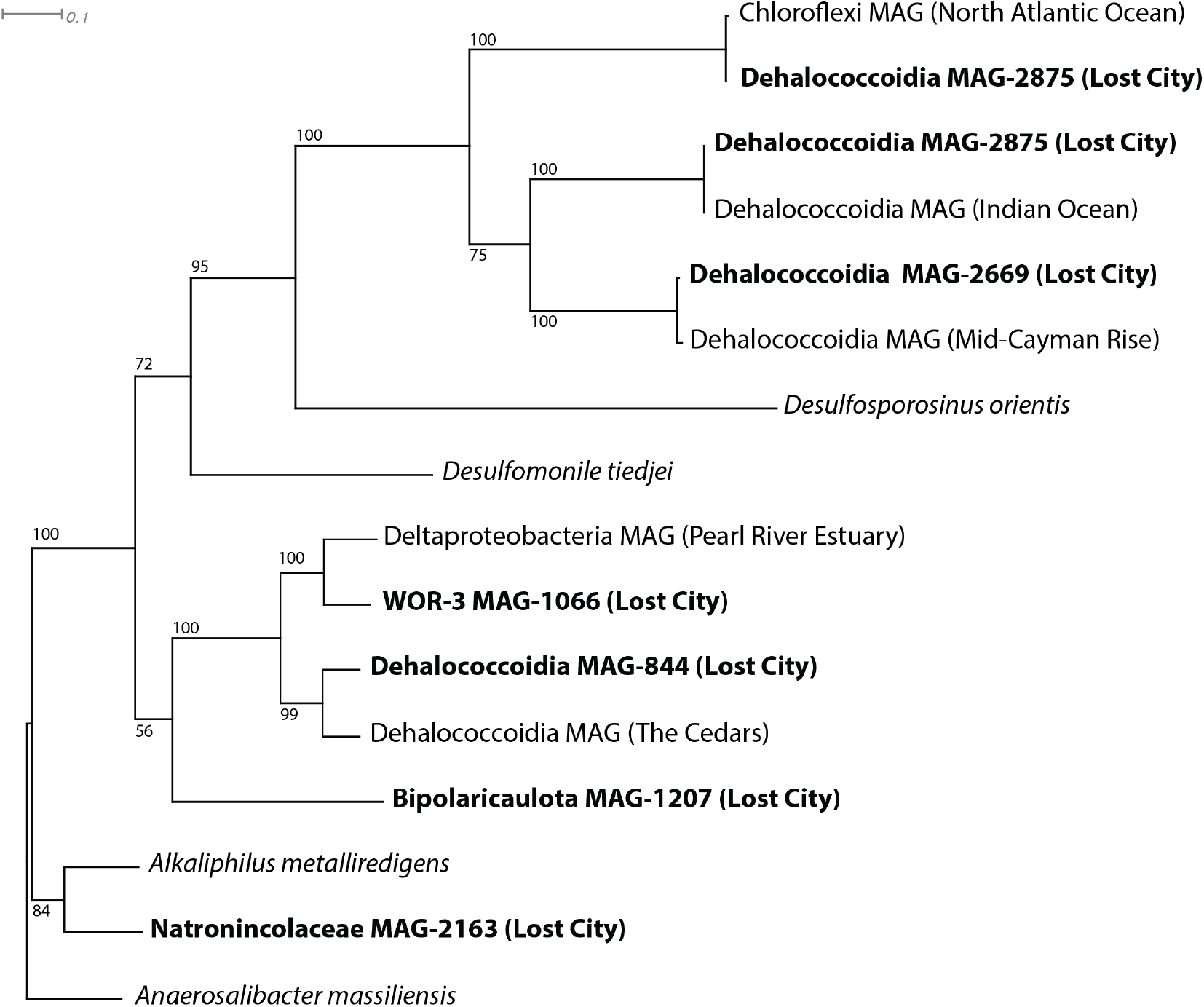
Phylogeny of GrdB (beta subunit of glycine reductase). Lost City Bipolaricaulota, Dehalococcoidia, WOR-3, and Natronincolaceae MAGs share moderate sequence similarity (58-88% amino acid identities) with sequences from other MAGs (including one from another site of serpentinization, The Cedars), but limited similarity with sequences from characterized species. Lost City Dehalococcoidia MAGs that belong to the SAR202 marine cluster, including two copies from MAG-2875, form a separate clade from other Lost City MAGs that are more likely to represent subseafloor organisms. A second Natronincolaceae MAG not shown here lacks GrdB but includes all other genes associated with glycine reductase (**Supplemental Table S5**). Bootstrap support values are shown for each node. Sequences and accession IDs are provided in the Zenodo-archived GitHub repository accessible via DOI: 10.5281/zenodo.5798015.

**Supplemental Figure S12.**
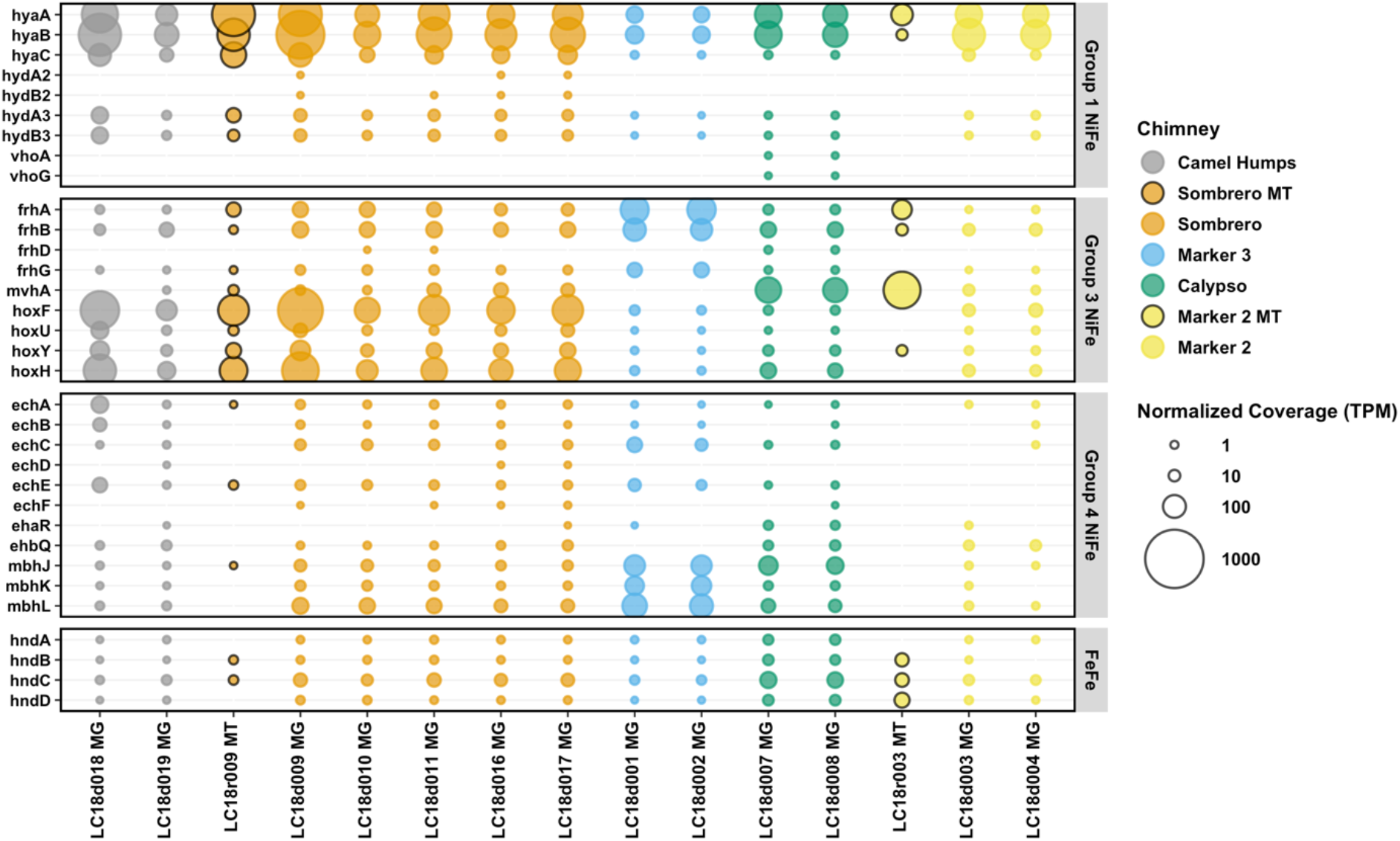
Abundance of predicted hydrogenase sequences in Lost City hydrothermal fluid samples. Metagenomic coverage was normalized to predicted protein length and to the size of the metagenome or metatranscriptome library. The final normalized coverage is reported as a proportional unit (transcripts/fragments per million; TPM) suitable for cross-sample comparisons. Bubbles representing coverage in metatranscriptomes (MT), rather than metagenomes (MG), are highlighted with black borders. Genes are defined with KEGG Orthology; see **Supplemental Table S5**.

**Supplemental Figure S13.**
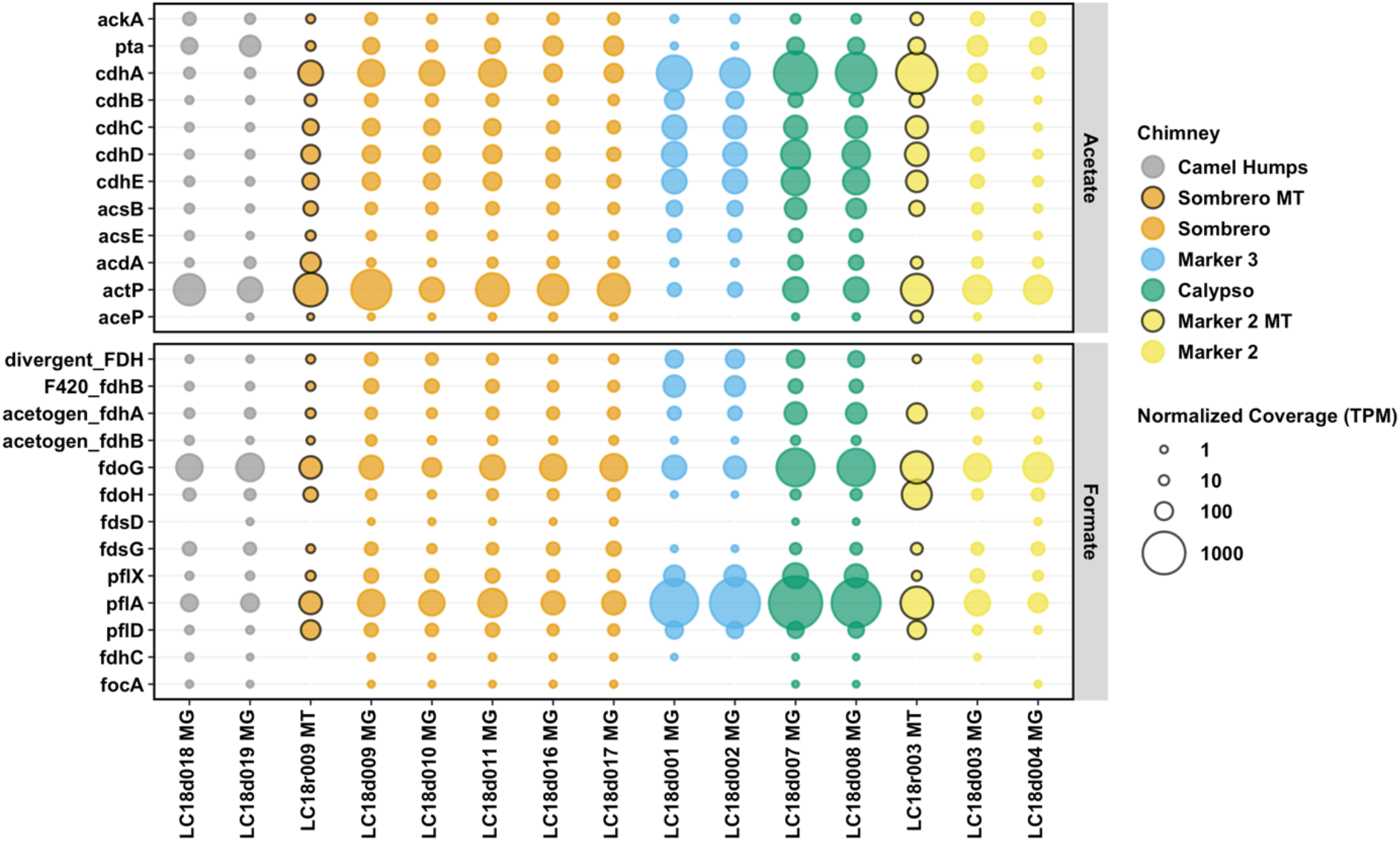
Abundances of predicted sequences associated with acetate and formate metabolism in Lost City hydrothermal fluid samples. Metagenomic coverage was normalized to predicted protein length and to the size of the metagenome or metatranscriptome library. The final normalized coverage is reported as a proportional unit (transcripts/fragments per million; TPM) suitable for cross-sample comparisons. Bubbles representing coverage in metatranscriptomes (MT), rather than metagenomes (MG), are highlighted with black borders. Genes are defined with KEGG Orthology; see **Supplemental Table S5**.

**Supplemental Figure S14.**
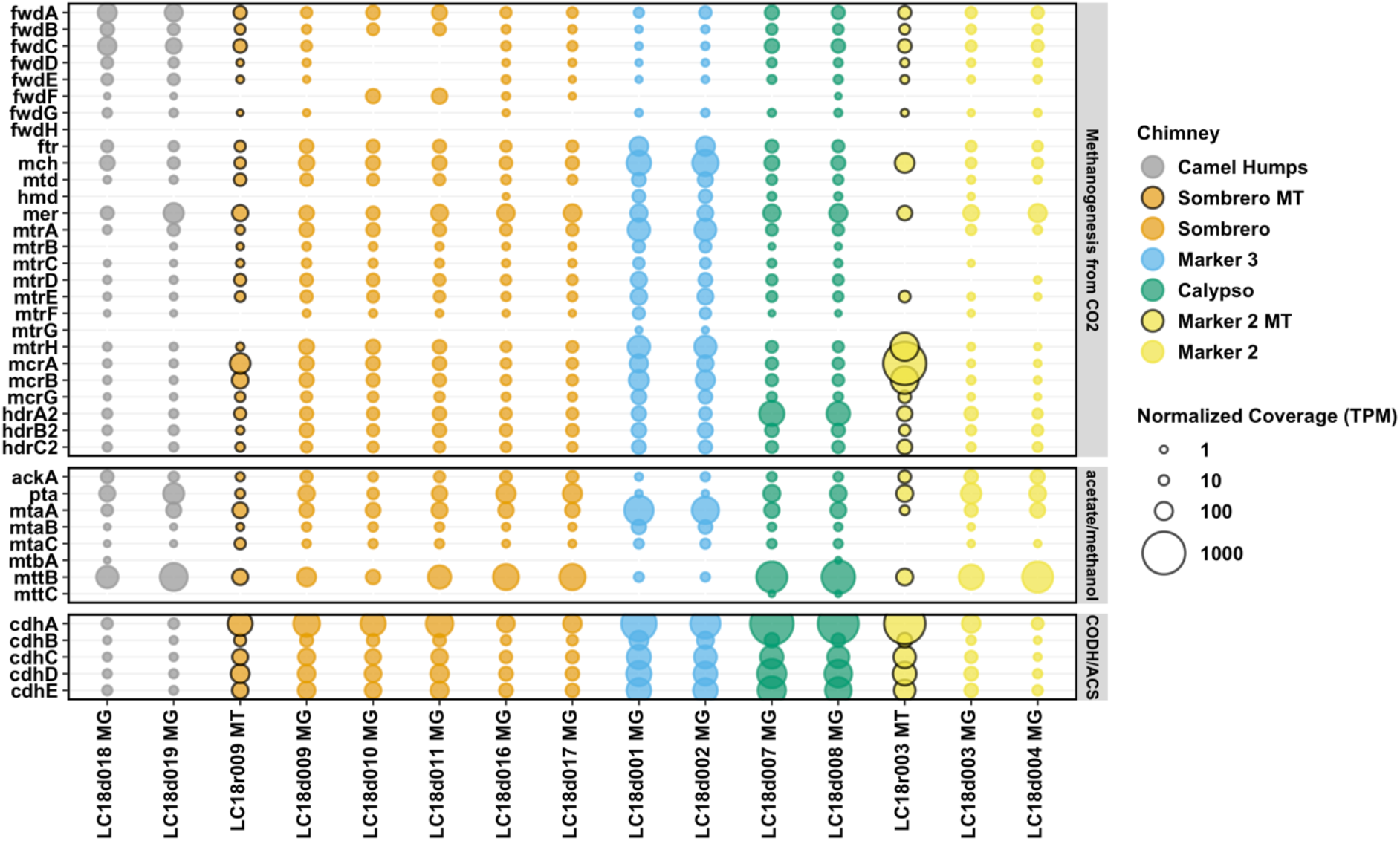
Abundance of predicted sequences associated with methanogenesis in Lost City hydrothermal fluid samples. Metagenomic coverage was normalized to predicted protein length and to the size of the metagenome or metatranscriptome library. The final normalized coverage is reported as a proportional unit (transcripts/fragments per million; TPM) suitable for cross-sample comparisons. Bubbles representing coverage in metatranscriptomes (MT), rather than metagenomes (MG), are highlighted with black borders. Genes are defined with KEGG Orthology; see **Supplemental Table S5**.

**Supplemental Figure S15.**
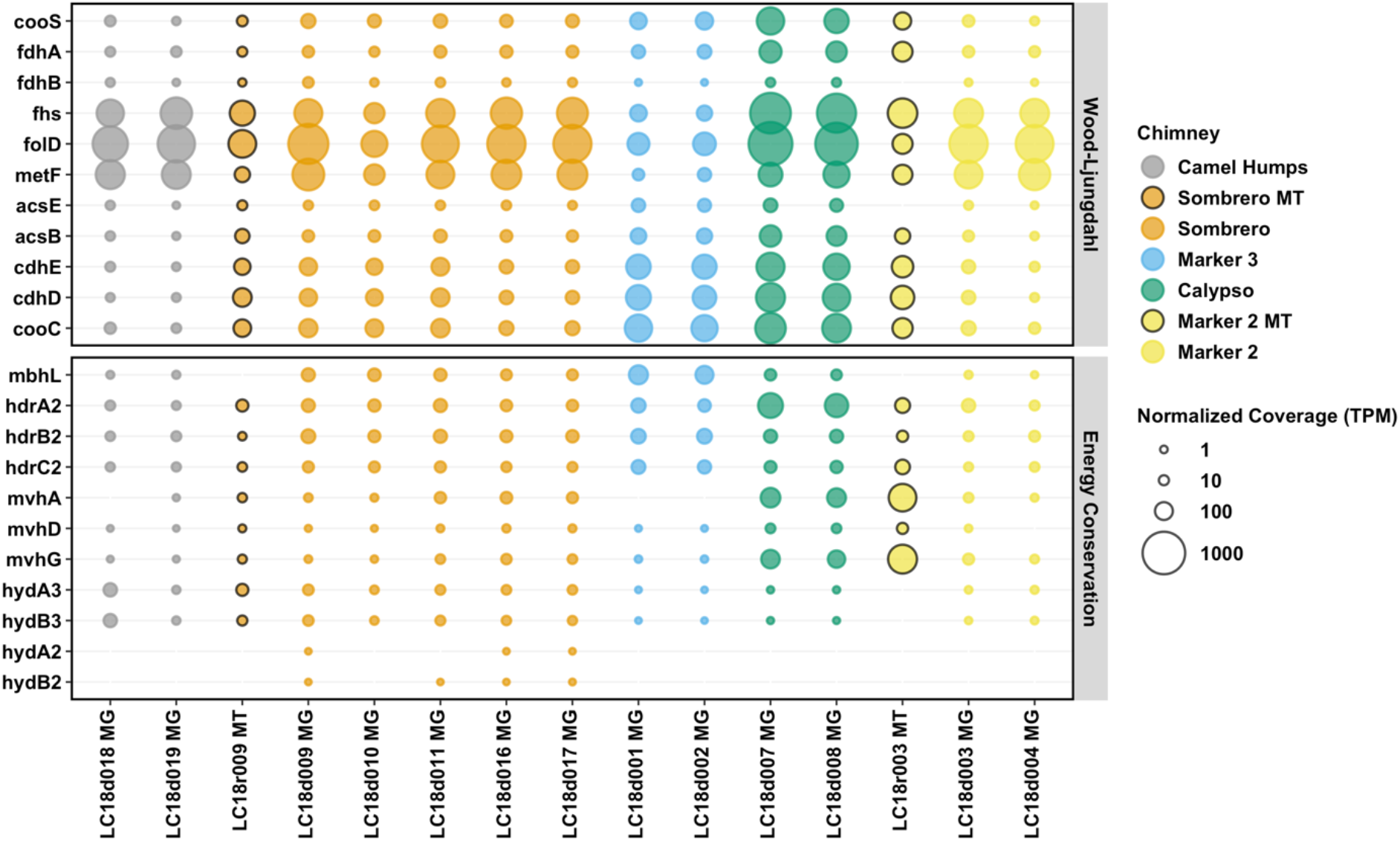
Abundance of predicted sequences associated with acetogenesis in Lost City hydrothermal fluid samples. Metagenomic coverage was normalized to predicted protein length and to the size of the metagenome or metatranscriptome library. The final normalized coverage is reported as a proportional unit (transcripts/fragments per million; TPM) suitable for cross-sample comparisons. Bubbles representing coverage in metatranscriptomes (MT), rather than metagenomes (MG), are highlighted with black borders. Genes are defined with KEGG Orthology; see **Supplemental Table S5**.

