## Supplemental Material for "Metabolic strategies shared by basement residents of the Lost City hydrothermal field"

### SUPPLEMENTAL METHODS

#### Collection of hydrothermal fluid samples

Hydrothermal fluid samples were collected from actively venting chimneys at the Lost City hydrothermal field (**Figure 1**) using ROV *Jason* during the 2018 Lost City expedition aboard R/V *Atlantis* (AT42-01). The present study included hydrothermal fluid samples from seven chimney locations: Camel Humps, Sombrero, Marker 3, Marker C, Calypso, Marker 2, and Marker 8 (**Table 1; Supplemental Table S1**). Several fluid samples collected from the Beehive chimney, where the highest fluid temperatures have been measured, yielded only low-quality DNA sequences and are not included in this study. The Sombrero site was sampled on two different ROV *Jason* dives, and samples from the separate dives are labeled as Sombrero1 or Sombrero2 when appropriate. Fluid samples collected from Markers C, 2, 3, and 8 were included in an early microbial diversity study (Brazelton et al., 2006), but microbial diversity data from the other chimneys are reported here for the first time.

On the seafloor, venting fluids were slowly pumped through 0.2 µm Millipore Sterivex cartridge filters or into acid-washed Kynar bags with the HOG sampler (Lang & Benitez-Nelson, 2021). Samples intended for RNA extractions were collected into 2 L Kynar bags containing 67 mL of a stop solution (97.5% 200 proof ethanol, 2.5% Trizol LS; Thermo Fisher). Fluid temperatures were monitored in real-time during sampling with a probe embedded into the sampler intake. Concentrations of sulfate, hydrogen sulfide, and magnesium were measured according to standard methods (Butterfield & Massoth, 1994). The estimated precision of the analyses is 3% for magnesium, 2% for sulfate, and 4% for hydrogen sulfide (all as relative standard deviation).

Chemical analyses were conducted on aliquots of the same fluid samples used for DNA and RNA sequencing. Separate chemical measurements were conducted on separate fluid samples dedicated for the chemistry analyses reported by (Aquino et al., In Revision).

Immediately upon shipboard recovery of ROV *Jason*, all Sterivex filters were stored at -80 ºC. Kynar bags were subsampled for shipboard aqueous chemistry measurements (pH, magnesium, sulfate, and sulfide) and filtered through Sterivex filters, which were subsequently frozen at -80 ºC. All frozen filters were shipped overnight to the University of Utah on dry ice or in liquid nitrogen vapor shippers.

**Cell counts**

Aliquots of all fluid samples were preserved in 3.7% formaldehyde and stored at 4 ºC. In the Anderson lab (Carleton College), preserved fluids were filtered onto 0.22 μm glass fiber filters and stained with DAPI (Porter & Feig, 1980). Cells were enumerated directly with an epifluorescence microscope, and a minimum of 20 fields of view were counted per sample. The median number of cells per field of view was recorded as the result for that sample, and the median value among all replicates is reported as the value for each location in **Table 1**.

**Bags supplemented with formate or methane**

On each dive, some of the 2 L Kynar bags were supplemented with 13C-enriched bicarbonate, formate, or methane (Cambrige Isotope Laboratories) to test for conversion of these compounds into dissolved inorganic carbon (DIC) or methane inside the Kynar bags. Thus, these Kynar bags served as seafloor incubation experiments. In addition, each bag contained 0.2 g of dithiothreitol, intended as a redox buffer. Upon shipboard recovery, the bags were subsampled and processed for DNA sequencing as described above, and additional aliquots of the fluid sample were collected for analysis of $^{13}C$ enrichment of methane and DIC. In the Lang lab (U.

South Carolina), the ratio of $^{13}C/^{12}C$ of headspace $CO_2$ was analyzed using a Thermo Scientific GasBench-Delta V Plus Isotope Ratio Mass Spectrometer (IRMS). The $^{13}C/^{12}C$ of $CH_4$ was analyzed with a Thermo Scientific Gas Chromatograph-IsolinkII-IRMS equipped with an Agilent GS - CarbonPlot column (30 m x 0.320 mm i.d., 1.50 um film thickness).

None of the bags supplemented with $^{13}C$-enriched formate or DIC produced methane with a $\delta^{13}C$ that was distinguishable from that of methane native to Lost City fluids (approximately -10 ‰) (Kelley et al., 2005; Proskurowski et al., 2008). Therefore, we conclude that no methane was produced from DIC or formate inside the Kynar bags.

Bags supplemented with $^{13}C$-enriched methane and containing fluids from Beehive, Sombrero, Calypso, Marker 8, and perhaps Marker 2 produced DIC with $\delta^{13}C$ values significantly greater than that of DIC native to Lost City fluids (< 1.5 ‰) (Kelley et al., 2005; Proskurowski et al., 2008). Furthermore, all of the bags supplemented with $^{13}C$-enriched formate produced DIC with enriched $\delta^{13}C$ values (**Supplemental Table S8**). Therefore, we conclude that methane and formate were oxidized to DIC in most of the Kynar sample bags.

Amplicon sequencing was conducted on each fluid sample separately, and no consistent differences in microbial community composition were detected among fluids collected in incubation bags compared to *in situ*-filtered Sterivex filters (**Supplemental Table S2**). Nevertheless, separate metagenome libraries were constructed with DNA prepared from *in situ*-filtered Sterivex filters and from incubation bags (**Supplemental Table S1**).

**Extraction and purification of DNA and RNA**

Extraction of DNA from all Sterivex filters (including *in situ*-filtered fluids and fluids collected in Kynar bags and filtered shipboard) was conducted as described previously (Brazelton et al.,

2017; Thornton et al., 2020). The full protocol is available in the Zenodo-archived GitHub repository (DOI: 10.5281/zenodo.5798015) and on protocols.io (DOI: dx.doi.org/10.17504/protocols.io.bykqpuvw). Cell lysis was performed at 65 ºC in a pH 10 extraction buffer followed by bead-beating with 0.1 mm glass beads. Lysates were purified with pH 8 phenol/chloroform/isoamyl alcohol extractions, precipitated overnight in ethanol, resuspended in low-EDTA TE, and further purified with magnetic beads (Rohland & Reich, 2012).

Total RNA was extracted from the Sterivex filters with a modification of the DNA extraction protocol optimized for RNA. The full protocol is available in the Zenodo-archived GitHub repository (DOI: 10.5281/zenodo.5798015) and on protocols.io (DOI: dx.doi.org/10.17504/protocols.io.bykspuwe). Cell lysis was performed at room temperature in a pH 7 extraction buffer, followed by bead-beating with 0.1 mm glass beads. Lysates were purified with pH 5.2 phenol/chloroform/isoamyl alcohol extractions, precipitated overnight in ethanol, resuspended in ultrapure water, and further purified with magnetic beads (Rohland & Reich, 2012). First-strand synthesis of cDNA was performed with SuperScript IV Reverse Transcriptase and random hexamers (Thermo Fisher).

**Sequencing of 16S rRNA amplicons**

Sequencing of amplicons generated from 16S rRNA genes and cDNA was performed at the Genomics Core Facility at Michigan State University on an Illumina MiSeq instrument using dual-indexed Illumina fusion primers targeting the V4 region of the 16S rRNA gene (Kozich et al., 2013). PCR products were batch-normalized and pooled using Invitrogen SequalPrep DNA Normalization plates. The pool was cleaned up using AmpureXP magnetic beads and quality-controlled and quantified using a combination of Qubit dsDNA HS, Agilent 4200 TapeStation HSDNA1000, and Kapa Illumina Library Quantification qPCR assays. This pool was loaded onto an Illumina MiSeq standard v2 flow cell, and sequencing was performed in a 2 x 250 bp paired-end format using a 500 cycle v2 reagent cartridge. Base calling was done by Illumina Real Time Analysis (RTA) v1.18.54, and the output of RTA was demultiplexed and converted to fastq format with Illumina Bcl2fastq v2.19.1.

**Analysis of 16S rRNA ASVs**

Amplicon sequence variants (ASVs) were inferred from 16S rRNA amplicon sequences with DADA2 v. 1.10.1 (Callahan et al., 2016) after removal of primer sequences with cutadapt v. 1.15 (Martin, 2011). Sequences from different sequencing runs were processed with DADA2 separately, and then the resulting ASVs from all sequencing runs were merged to form a final, non-redundant count table of ASVs. Potential contaminants were removed with the decontam package (Davis et al., 2018) using both the "frequency" and "prevalence" modes. In "frequency" mode, ASVs were removed if their counts were significantly correlated ($p < 0.02$) with DNA concentration (as measured by Qubit fluorometric quantification, Thermo Fisher). In "prevalence" mode, ASVs were removed if they were significantly more likely ($p < 0.02$) to be present in one of three likely contamination sources (ambient lab air, surface seawater, or DNA extraction blanks) than in any sample of hydrothermal fluid. Samples of ambient air in the R/V *Atlantis* shipboard laboratory and our laboratory at the University of Utah were obtained as previously described (Motamedi et al., 2020). Samples of surface seawater were obtained during a previous study at the same location above the Atlantis Massif (Motamedi et al., 2020) and also from the shipboard laboratory "tap" water produced from surface seawater by the R/V *Atlantis*. DNA extraction blanks were obtained by Motamedi et al. by subjecting sterile Sterivex filters to the DNA extraction protocol described above. All contamination control samples were sequenced on the same sequencing runs as described above for the hydrothermal fluid samples. A total of 1,823 ASVs (9% of all ASVs) representing 109,824 sequence counts (2% of all sequence counts) were removed from downstream analyses by decontam. These removed ASVs are provided in **Supplemental Table S2**.

Taxonomic classification of all ASVs was performed with DADA2 using the SILVA reference alignment (SSURefv132) and taxonomy outline (Quast et al., 2013; Yilmaz et al., 2014). Bubble plots of ASVs were drawn with ggplot2 using proportional abundances. The ordination plot was generated with phyloseq v1.26.1 (McMurdie & Holmes, 2013) using an unconstrained NMDS ordination of Morisita-Horn dissimiliarity values. The stress of the fit to two dimensions was 0.05 after 20 tries without a convergent solution. Very similar results were produced with MDS and CCA ordinations and with Bray-Curtis dissimilarity values.

Differential abundances of ASVs between the Marker 3 and Camel Humps locations were calculated with DESeq2 (Love, Huber & Anders, 2014) as implemented by phyloseq (McMurdie & Holmes, 2013). ASVs with variance $<10^{-5}$ were filtered out prior to the test, and those ASVs with an adjusted p-value $<0.05$ were considered to be differentially abundant.

**Sequencing of metagenome libraries**

Metagenome libraries were constructed with size-selected, sonicated DNA fragments of 500-700 bp with the NEBnext Ultra DNA II library kit for Illumina (E7645S), as previously described (Thornton et al., 2020). Paired-end sequencing (2 x 125 bp) of metagenomic libraries was conducted at the University of Utah High-Throughput Genomics Core Facility at the Huntsman Cancer Institute with an Illumina HiSeq2500 platform. Sequencing libraries (25 pM) were chemically denatured and applied to an Illumina HiSeq v4 paired end flow cell using an Illumina cBot.  Hybridized molecules were clonally amplified and annealed to sequencing primers with reagents from an Illumina HiSeq PE Cluster Kit v4-cBot. Following transfer of the flowcell to an

Illumina HiSeq 2500 instrument (HCS v2.2.38 and RTA v1.18.61), a 125 cycle paired-end sequence run was performed using HiSeq SBS Kit v4 sequencing reagents (FC-401-4003).

**Sequencing of metatranscriptome libraries**

Only two samples (one from Marker 2 and one from Sombrero) contained sufficient total RNA to attempt metatranscriptome sequencing. The two metatranscriptome libraries were constructed and sequenced by the University of Utah High-Throughput Genomics Core Facility at the Huntsman Cancer Institute. Total RNA was hybridized with NEBNext rRNA Depletion Solution Bacteria (E7850L) to substantially diminish rRNA from the samples. Stranded RNA sequencing libraries were prepared using the NEBNext Ultra II RNA Library Prep Kit for Illumina (E7770L). Purified libraries were qualified on an Agilent Technologies 2200 TapeStation using a D1000 ScreenTape assay. The molarity of adapter-modified molecules was defined by quantitative PCR using the Kapa Biosystems Kapa Library Quant Kit. Individual libraries were normalized to 10 nM, and equal volumes were pooled. Sequencing libraries were chemically denatured and applied to an Illumina NovaSeq flow cell using the NovaSeq XP workflow (20043131). Following the transfer of the flowcell to an Illumina NovaSeq 6000 instrument, a 150 cycle paired-end sequence run was performed using a NovaSeq 6000 S4 reagent Kit v1.5 (20028312).

**Quality control and taxonomic classification of metagenome and metatranscriptome sequences**

Demultiplexing and conversion of the raw sequencing base-call data were performed through the CASAVA v1.8 pipeline. Adapter sequences and PhiX were removed from all reads with BBDuk (part of the BBTools suite, v35.85 (Bushnell, Rood & Singer, 2017)). Quality trimming was performed with our seq-qc package (https://github.com/Brazelton-Lab/seq-qc) as previously described (Thornton et al., 2020). Each library yielded 22-192 million reads (for a total of 1.4 billion reads among all metagenome libraries) after these quality control steps, representing 58-

82% of the number of raw reads. The two metatranscriptome libraries yielded 249 million reads (Marker 2) and 467 million reads (Sombrero) after quality filtering, representing 82% and 88%, respectively, of the original raw reads. Ribosomal RNA sequences were identified in the two metatranscriptomes with SortMeRNA (Kopylova, Noé & Touzet, 2012), resulting in the removal of 77% of reads from the Marker 2 metatranscriptome and 90% of reads from the Sombrero metatranscriptome. Quality-filtered, unassembled reads were assigned taxonomy with Kaiju and the NCBI nr+euk database (Menzel, Ng & Krogh, 2016). Kaiju was unable to classify 46-86% of reads, with the lowest percentage of unclassified reads in metatranscriptomes and the highest percentage of unclassified reads in the two Marker 3 metagenomes and one Sombrero metagenome (**Supplemental Table S3**). An interactive Krona plot is provided in the Zenodo-archived GitHub repository (DOI: 10.5281/zenodo.5798015).

**Metagenomic assembly**

Sequences from metagenomic libraries were assembled with Megahit v1.1.1 (Li et al., 2016), using kmers of 27 to 127. A pooled, "all fluids" Megahit assembly was performed with metagenomic reads from all 13 libraries (representing 37 fluid samples collected in Sterivex filters and Kynar bags; see **Supplemental Table S1**). Genes were predicted with Prodigal v2.6.3 (Hyatt et al., 2010) in meta mode. Predicted protein sequences were queried against the KEGG release 83.2 prokaryotes database with Diamond v0.9.14 (Buchfink, Reuter & Drost, 2021).

In addition to the "all fluids" Megahit assembly, chimney-specific assemblies were conducted with metaSPAdes v3.13.0 (Nurk et al., 2017) as implemented by the KBase platform (kb_SPAdes v.1.2.4) (Arkin et al., 2018). A chimney-specific assembly with reads obtained from Marker 3 was performed with Megahit v1.1.1 because metaSPAdes repeatedly failed when it attempted to assemble Marker 3 reads. An overview of the metagenomic analysis workflow is illustrated in **Supplemental Figure S3**.

**Binning of Metagenome-Assembled Genomes (MAGs)**

Binning of metagenome-assembled genomes (MAGs) from the “all fluids” and Marker 3 Megahit assemblies was conducted with BinSanity using the Binsanity-lc workflow v0.2.6.2 and a minimum contig size of 3 kb (Graham, Heidelberg & Tully, 2017). Binning of MAGs from the chimney-specific metaSPAdes assemblies was conducted with MaxBin2 v2.2.4 (kb_maxbin v.1.1.1) (Wu, Simmons & Singer, 2016), MetaBAT2 v2.2 (kb_metabat v.2.3.0) (Kang et al., 2019), and DAS Tool v1.1.2 (kb_das_tool v.1.0.6) (Sieber et al., 2018). The ANME-1 MAG was reconstructed from a Megahit assembly of Calypso reads with MetaBAT2 and DAS Tool. MAGs were assigned taxonomic classifications with GTDB-Tk v1.5.1 (reference data version r202;(Chaumeil et al., 2020)). Individual contigs were assigned taxonomic classifications with MMseqs2 (Mirdita et al., 2021). Gene products were predicted by annotation with Prokka v1.14.5 and its default databases (Seemann, 2014) with further function prediction provided by GhostKOALA v2.2 (Kanehisa, Sato & Morishima, 2016). Protein identifications and predicted functions were supplemented by results from InterProscan 5 (v5.52-86.0) (Jones et al., 2014), FeGenie (Garber et al., 2020), and dbCAN2 (Zhang et al., 2018). Completeness and contamination of initial automated MAGs were estimated with CheckM v1.0.5 (Parks et al., 2015), and the completeness and redundancy of the final refined MAGs were estimated with anvi’o v7 (Eren et al., 2021). Selected MAGs were refined by re-assembly with metaSPAdes v3.11.1 using reads mapped to MAGs and contigs with matching taxonomic classifications, followed by manual curation in anvi'o v7 (Eren et al., 2015) according to the differential coverage of contigs among samples. The final, refined MAGs represent 0.02 – 0.84% of the total read coverage in each metagenome or metatranscriptome library.

**Coverages of genes, contigs, and MAGs**

Quality-filtered metagenome and metatranscriptome sequences were mapped to the “all fluids” Megahit assembly with Bowtie2 v2.3.2 (Langmead & Salzberg, 2012). Bowtie2 mapping rates to the “all fluids” assembly were 67-71% for Camel Humps, 82-89% for Sombrero1, 60-71% for Sombrero2, 95% for Marker 3, 67-74% for Calypso, and 56-60% for Marker 2. Bowtie2 mapping rates to the “all fluids” assembly were 21% before rRNA removal and 14% after rRNA removal for the Marker 2 metatranscriptome and 83% before rRNA removal and 74% after rRNA removal for the Sombrero1 metatranscriptome. Artificial duplicates were removed from the bam files using the MarkDuplicates function of Picard Tools v2.17.8 (*Picard Toolkit*, 2019).

The coverage for each predicted protein was calculated as transcripts (or fragments) per million (TPM) with count_features v1.3.0, part of our seq-annot package (Thornton et al., 2020). TPM is a proportional unit (multiplied by one million for convenience) that is normalized to the length of each predicted protein sequence as well as to the total library size. Coverages of both metagenomes and metatranscriptomes were calculated in the same way and reported in the same units (TPM), although the metagenomic coverages reflect fragments instead of transcripts (Thornton et al., 2020).

The coverage of each MAG was calculated as the weighted sum of the normalized, proportional coverages (in TPM) of its member contigs. The contig coverages were obtained by mapping all unassembled reads against the re-assembled, refined MAGs with Bowtie2 and then calculating the average coverage per position of each contig with the genomecov command (with the option -pc) in bedtools v2.30.0 (Quinlan & Hall, 2010). Normalized coverages in units of TPM were calculated by dividing the average coverage per position by the total number of read pairs for that library.

**Phylogenetic Analyses**

Phylogenetic trees of 16S rRNA genes were constructed with alignments obtained from the SILVA alignment server (Pruesse, Peplies & Glöckner, 2012) and RAxML v8.2.11 (Stamatakis, 2014) using the gamma model of rate heterogeneity and the "-f a" option to construct 100 rapid bootstrap searches and 20 maximum likelihood searches. The *Thermodesulfovibrionales* ASV from the Samail Ophiolite, Oman (Rempfert et al., 2017) was generated with DADA2 as described above with reads accessed via SRA accession SRR5000240. Phylogenies of McrA, GrdB, and carbonic anhydrase were constructed with Clustal Omega alignments (Madeira et al., 2019) and RAxML as described above but with automated selection of the rate model. All sequences and accessions are provided in the Zenodo-archived GitHub repository accessible via DOI: 10.5281/zenodo.5798015.

---

### SUPPLEMENTAL RESULTS:

#### *DETAILED COMPARISONS OF HYDROTHERMAL FLUID SAMPLES*

**Markers 3 and C**

The strong similarity of community compositions from Markers 3 and C is remarkable considering that fluids from Marker 3 were sampled at much lower temperatures (<20 ºC) than those from Marker C (up to 80 ºC). Temperatures exceeding 55 ºC were measured at Marker 3 during the same dive, but the temperatures of the fluid samples (i.e. measured in-line, during sampling) used for DNA-based analyses were lower, most likely due to cooling of the fluids as they exit the chimney. In addition, the pH, sulfate, sulfide, and magnesium levels of Marker 3 fluids are more similar to those of seawater, compared to Marker C fluids. These results suggest that the cooler fluids venting from Marker 3 experience more dilution with ambient seawater compared with the warmer fluids venting from Marker C, but that they share a common subsurface source, which may explain the similar microbial compositions.

Marker 3 fluids were rich in metagenomic sequences classified as family *Methanosarcinaceae,* which includes the dominant archaeal phylotype previously detected in Lost City chimneys (Schrenk et al., 2004; Brazelton et al., 2010, 2011). Archaeal sequences were much more abundant in metagenomes than in the 16S rRNA amplicon libraries. For example, *Methanosarcinaceae* is the most abundant family in Marker 3 metagenomic reads (1-2% of all reads and 20-21% of all reads that could be classified to the family level by Kaiju; **Supplemental Table S3**), even though *Methanosarcinaceae* ASVs have lower relative abundance than several bacterial ASVs in the same samples, suggesting a bias against archaeal sequences in the ASV dataset.

Venting fluids collected near Markers C and 3 were particularly enriched in taxa representing potential sulfate-reducing bacteria (SRB). *Thermodesulfovibrionia* and *Desulfotomaculum* ASVs dominated these fluids, but they were much less abundant in the RNA fraction of the Marker C sample (**Figure 3**). ASVs classified as *Desulfocapsa* were abundant in Marker 3 fluids, present in the RNA fraction from Marker C, and absent in the DNA fraction from Marker C. *Desulfobulbus* ASVs were absent in both Markers C and 3 except for the RNA fraction from Marker C, and they were prevalent in Sombrero fluid samples. *Desulfobulbus* and *Desulfocapsa* both belong to the family *Desulfobulbaceae*, which had 30-fold less coverage in Marker 3 metagenomes than *Methanosarcinaceae* and 8-fold less coverage than the *Nitrospiraceae* family that includes *Thermodesulfovibrio* (**Supplemental Table S3**).

Marker 3 metagenomes were also distinctive in their high proportions of Candidatus Patescibacteria (**Supplemental Figure S2**). Patescibacteria were rare in all samples except for Marker 3 fluids, where they represented 33-36% of all classified sequences (**Supplemental Table S3**). Hundreds of ASVs were classified as Patescibacteria (predominantly Paceibacteria and Gracilibacteria), but they did not represent a large fraction of the counts in any of the fluid samples (**Supplemental Table S2**).

**Camel Humps and Sombrero**

Fluids venting from Camel Humps contained a remarkably even distribution of ASVs that included *Sulfurovum*, *Sulfurospirillum, and Thiomicrorhabdus* at similar abundances as taxa typically associated with ambient seawater (e.g. *Alteromonas*, *Roseobacter, Halomonas*). This community composition strongly contrasts with that of Marker 3, even though both locations are at the summit of the central Poseidon structure (**Figure 1**). Of the 100 ASVs most common in Camel Humps fluids, 70 of these were significantly less abundant in Marker 3 fluids (**Supplemental Table S2**). The differences between Camel Humps and Marker 3 cannot be explained by mixing of the same hydrothermal fluid with ambient seawater because the bacteria at Camel Humps belong to lineages associated with hydrothermal environments (e.g., *Sulfurovum*, *Sulfurosprillum*, *Thiomicrorhabdus*), not ambient seawater. Moreover, the temperatures measured during the sampling of Camel Humps fluids were higher than those of Marker 3 fluids, which is the opposite of what would be expected if dilution by seawater had been responsible for the observed trends. Instead, these results suggest a difference in the subsurface source of the hydrothermal fluids venting from these locations. Although Marker 3 and Camel Humps are located next to each other, they are visually distinct chimney structures that could potentially host fluids venting from distinct subsurface sources (**Supplemental Figure S2**). Alternatively, the distinctive assemblage of taxa at Camel Humps may represent biofilm communities that were flushed into nearby venting fluids, swamping the less dense organisms derived from the subsurface. The two explanations are not mutually exclusive, and both indicate that the fluids collected from Camel Humps during this study are not representative of the subseafloor.

The Sombrero chimney is located on a ridge stretching from the main vent field toward the eastern wall of the Atlantis Massif (**Figure 1**), and it was sampled on two separate ROV *Jason* dives, with the second dive (Sombrero2) collecting fluids with higher temperatures and lower sulfate concentrations than the first dive (Sombrero1) (**Table 1**). Despite these different measurements at the time of sampling, the overall microbial composition was remarkably consistent across all Sombrero samples ranging in temperature from 10 – 74 ºC (**Figure 1**). Minor differences are nevertheless visible (**Figure 2**); for example, *Thiomicrorhabdus* and *Sulfurospirillum* dominated Sombrero fluids with lower temperature, higher sulfate, and lower sulfide. Warmer and more sulfidic Sombrero fluids included greater proportions of taxa that were also abundant in fluids from Markers 3 and C (**Figure 2**).

**Markers 2 and 8**

The most sulfidic fluids were collected from chimneys near Marker 2 and Marker 8, which are located on the western edge of the Lost City hydrothermal field (**Table 1**; **Figure 1; Supplemental Figure S1**), and their microbial compositions were distinct from all other fluids. One of the fluid samples from Marker 2 was dominated by a single ASV identical to the 16S rRNA gene of *Alteromonas macleodi* (**Supplemental Table S2**), a ubiquitous marine bacterium (Koch et al., 2020), suggesting substantial dilution of the sample with ambient seawater. Other samples from Marker 2 also contained bacteria that have been previously associated with sulfur oxidation in Lost City chimney biofilms, including the genera *Sulfurovum*, *Sulfurospirillum*, and *Thiomicrorhabdus* (Brazelton et al., 2010; Brazelton & Baross, 2010; McGonigle, Lang & Brazelton, 2020). Therefore, they are likely to be adapted to chimney habitats where sulfidic hydrothermal fluids are mixing with oxic seawater, and they are probably not representative of microbial communities inhabiting anoxic, subsurface environments. The ribosomal RNA fraction of Marker 2 fluids contained elevated relative abundances of a *Sulfurospirillum* ASV and a *Thiomicrorhabdus* ASV (**Figure 2**). Microbial taxa detected in fluids collected near Marker 8 were broadly similar to those of Marker 2 fluids, except that they were dominated by one *Sulfurovum* ASV that was rare in Marker 2 fluids (**Figure 2**).

**Calypso**

The Calypso chimney sits on the eastern wall of the Atlantis Massif, approximately 75 m from the large Poseidon edifice that dominates the Lost City field (**Figure 1**). The fluids venting from Calypso had higher sulfide concentrations than the fluids from Markers 3 and C, but less sulfide than in Markers 2 and 8 (**Table 1**). The overall microbial community structure resembled that of Sombrero fluids (**Figure 1**), although the ASVs with the highest counts in Calypso were also common in Markers 3 and C (**Figure 2**). The top *Thermodesulfovibrionia* ASV in Calypso fluids differed by one base from the ASV that dominated the fluids from Markers C and 3. This ASV was further enriched in the RNA fraction from Calypso, suggesting that it could have been metabolically active prior to sampling.

ASVs classified as the ANME-1b group of archaeal methanotrophs were most abundant in fluids from Calypso (**Figure 2**). ANME-1 sequences were rare or absent in almost all other fluid samples except for a few samples from Markers C and 3 and the RNA (but not DNA) fraction from one sample of fluids at Marker 2. These results are consistent with our previous studies that documented very high relative abundances of ANME-1 in cooler chimneys at the periphery of the field and trace levels of ANME-1 DNA in hot chimneys of the central Poseidon complex (Brazelton et al., 2006, 2010).

Calypso fluids were rich in Chloroflexi ASVs and metagenomic sequences primarily classified as class *Dehalococcoidia* (**Supplemental Tables S2-S3)**, but ASVs classified as *Anaerolineae*, TK10, and KD4-96 were also present at low levels. The Chloroflexi MAG from Lost City biofilms previously reported in (McGonigle, Lang & Brazelton, 2020) is a member of the *Anaerolineae*.

*Dehalococcoidia* ASVs were generally most abundant in Calypso fluids, but most ASVs were broadly distributed (**Figure 2**).

---

**SUPPLEMENTAL RESULTS:**

### ***DETAILED DESCRIPTIONS OF METAGENOME-ASSEMBLED GENOMES (MAGs)***

#### ***Methanosarcinaceae* MAG**

The predominant *Methanosarcinaceae* ASV was classified as genus *Methanosalsum* in the SILVA database, but our phylogenies of 16S rRNA and mcrA indicate that the Lost City *Methanosarcinaceae* are not monophyletic with any previously characterized genera (**Supplemental Figure S5**). Over 254 bases, the *Methanosarcinaceae* ASV is 94% similar to the 16S rRNA gene of *Methanococcoides methylutens* and 95.7% similar to that of *Methanohalophilus mahii*.

The *Methanosarcinaceae* MAG contained 1276 coding sequences (CDS) and was estimated to be 84% complete with 1% redundancy. It shares 99% average nucleotide identity (ANI) with the MAG we previously recovered from a Lost City biofilm (McGonigle, Lang & Brazelton, 2020). It is most abundant in Marker 3 fluids (**Figure 3**), and it is more abundant in those Sombrero fluids that show more mixing with seawater (lower temperature, more sulfate, less sulfide), consistent with the distribution of *Methanosarcinaceae* ASVs (**Figure 2**).

*Methanosarcinaceae* MAG-1276 has a remarkably low GC content (29%), which may help explain why the first metagenomic studies of Lost City biofilms recovered surprisingly few archaeal sequences (Brazelton & Baross, 2009, 2010). It encodes the core pathway for methanogenesis from carbon dioxide (**Supplemental Table S5**). It contains predicted sequences for $F_{420}$-reducing hydrogenase (FrhAB), which is required by all methanogens that reduce carbon dioxide with $H_2$ (Mand & Metcalf, 2019). In addition, two genes predicted to encode subunits of Ech hydrogenase (EchCE) are present. FrhB and EchCE are located on the same contig as a multicomponent $Na^+$:$H^+$ antiporter (MrpACBD) and the MAG's only gene annotated as a subunit of NADH-quinone oxidoreductase (NuoH).

The MAG encodes AMP-forming acetyl-CoA synthetase, as previously reported for *Methanosarcinaceae* in Lost City biofilms (McGonigle, Lang & Brazelton, 2020), which may enable acetoclastic methanogenesis. It also encodes MtaA and MtaC (**Supplemental Table S5**), two of the three proteins that enable the use of methanol as a substrate for methanogenesis in some methanogens.

*Methanosarcinaceae* MAG-1276 contains a complete 14-gene cluster (mbhA-N) encoding membrane-bound hydrogenase (**Figure 5; Supplemental Figure S8**). The same gene cluster, with conserved synteny, is also found in methanogens belonging to the order *Methanomicrobiales* (e.g. *Methanospirillum hungatei*) and in heterotrophs of the order *Thermococcales* (e.g. *Thermococcus kodakarensis*) (Thauer et al., 2010). The mbhL subunits from these methanogens have only 42-45% identities with the Lost City mbhL sequences reported here, which have greater similarly (~49% identities) to mbhL sequences from *Thermococcus*.

As we reported previously for chimney biofilms (Lang et al., 2018), the *Methanosarcinaceae* MAG encodes a formate dehydrogenase (FDH) that is similar to that of *Methanolobus* species, which are unable to use formate as a carbon source. The contig encoding this FDH also encodes one subunit of $F_{420}$-non-reducing hydrogenase (MvhD) and heterodisulfide reductase (HdrABC).

We identified additional, divergent FDH-like sequences associated with *Methanosarcinaceae*, ANME-1, *Thermodesulfovibrionales*, and Bipolaricaulota (**Figure 6; Supplemental Figure S9**). These divergent FDH-like sequences have not been previously described, though they are present in a few published genomes. The *Methanosarcinaceae* FDH-like sequence (**Figure 5**) shares 63% amino acid identities with a predicted protein in the *Desulfitibacter alkalitolerans* genome. In a previous study, we reported these divergent FDH-like sequences as bacterial based on their similarity to *Desulfitibacter* sequences (Lang et al., 2018). Although *Desulfitibacter alkalitolerans* can use formate as an electron donor, it encodes two additional homologs of fdhA that are not present in any Lost City metagenomes. A similar FDH-like sequence is encoded by *Dethiobacter alkaliphilus*, which is unable to grow on formate as its sole carbon source (Sorokin et al., 2008). The phylogeny in **Figure 5** is rooted with FdhF (anaerobic formate hydrogen lyase) from *Methanothermobacter thermautotrophicus* and *Methanothermobacter marburgensis.* These two species are unable to grow on formate (Kaster et al., 2011), and they lack a separate fdhABC operon that is found in *M. thermautotrophicus* st. CaT2, which can grow on formate.

No known transporters for formate were identified in this MAG, but one gene was predicted to encode a member of the oxalate:formate antiporter family. In *Oxalobacter formigenes*, this transporter enables uptake of oxalate with export of formate (Abe, Ruan & Maloney, 1996). Its presence was also observed in a formate-utilizing methanogen, which lacks any other formate transporters, in hyperalkaline groundwaters of the Samail Ophiolite (Fones et al., 2021). Potentially homologous oxalate:formate antiporters were also identified in two NPL-UPA2 MAGs and two *Natronincolaceae* MAGs from Lost City fluids. The four bacterial sequences shared 59-63% amino acid identities with the *Methanosarcinaceae* sequence.

#### *Methanocellales* MAG

A MAG classified as order *Methanocellales* (838 CDS; 84% complete with 1% redundancy) was only present in Calypso fluids (**Figure 3**). Curiously, no ASVs were classified as *Methanocellales*, and no previous studies have identified any methanogen taxa in Lost City samples other than *Methanosarcinaceae*.

*Methanocellales* MAG-838 encodes the key enzyme CODH/ACS, but it has an incomplete pathway for methanogenesis, including only four of the first five steps (FwdABCDFG, Ftr, Mtd, and Mer) and lacking all other steps, including the proteins required for methane production (methyl-coenzyme M reductase and heterodisulfide reductase) (**Supplemental Table S5**). Evidence for acetate utilization includes a predicted sequence for AMP-forming acetyl-CoA synthetase, which shares 73% amino acid identities with the homolog in the *Methanosarcinaceae* MAG, and a cation/acetate symporter (ActP). The MAG includes one [NiFe] hydrogenase and the Rnf complex (RnfCDGEAB), an energy-conserving ferredoxin:$NAD^+$-oxidoreductase (Biegel et al., 2011).

#### ANME-1 MAG

A MAG classified as ANME-1 (1099 CDS; 88% complete with 4% redundancy) was most abundant in Calypso fluids as well as the Marker 2 metatranscriptome, even though ANME-1 was very rare in the Marker 2 metagenomes. As expected for ANME-1 archaea, the Lost City ANME-1 MAG contains the core methanogenic pathway (**Supplemental Table S5**). The absence of cytochromes and presence of hydrogenases in this MAG was noted by (Chadwick et al., 2021) as consistent with the genomic features of the so-called “freshwater” clade of ANME-1, for which the genus “Candidatus Methanoalium” was proposed. One of the shared features within this clade, including the Lost City ANME-1 MAG, is a novel HdrABC-MvhADG complex, which is involved in the transfer of electrons derived from $H_2$ in methanogens.

The Lost City ANME-1 MAG contains the core methanogenic pathway (**Supplemental Table S5**), including $F_{420}$-dependent methylenetetrahydromethanopterin reductase (Mer). This gene is required for methanogenesis from carbon dioxide, but it is typically absent in ANME genomes, with at least one exception previously reported (Beulig et al., 2019). The MAG lacks all but one of the subunits of $N^5$-methyl-$H_4$MPT:coenzyme M methyltransferase (Mtr), which catalyzes the penultimate step of methanogenesis (and putatively the second step of reverse methanogenesis). It is present in most but not all ANME-1 genomes (Chadwick et al., 2021).

The ANME-1 MAG-1 also encodes the complete mbhA-N gene cluster for membrane-bound hydrogenase (**Figure 5; Supplemental Figure S8**), and each predicted gene in the cluster has the greatest similarity to the homolog in the *Methanosarcinaceae* MAG than to any other sequences in public databases. It lacks any established genes for FDH, but it includes a divergent FDH-like sequence that appears to be homologous to those found in other Lost City MAGs (**Figure 6; Supplemental Figure S9**). Distant homologs of FDH have been previously identified in ANME genomes, but they do not share significant sequence similarity with the FDH-like sequences reported here.

**Bipolaricaulota MAGs**

ASVs classified as Bipolaricaulota (named Acetothermia in the SILVA database) clustered into three distinct clades (**Supplemental Figure S6**) that correspond to the taxonomic classifications of three distinct Bipolaricaulota MAGs. The most abundant of the three Bipolaricaulota MAGs (1207 CDS; 86% complete with 0% redundancy) was classified by GTDB as species UBA7950 within class Bipolaricaulia, and it shares 99% ANI with the GTDB reference genome, which was assembled by (Parks et al., 2021) with sequences from a previously published Lost City biofilm metagenome (Lang et al., 2018). Bipolaricaulota MAG-1207 is most abundant in Sombrero and

Calypso, especially the Sombrero metatranscriptome (**Figure 3**). It encodes a nearly complete glycolysis pathway, an incomplete TCA cycle, an incomplete Wood-Ljungdahl pathway, the Rnf complex, and pyruvate formate-lyase (PflD) (**Supplemental Table S5**).

A second Bipolaricaulota MAG (1260 CDS; 89% complete with 1% redundancy) was classified as family UBA9294 within order UBA7950, and it shares only 77% ANI with the closest reference in GTDB (**Supplemental Table S4**). Bipolaricaulota MAG-1260 has a similar complement of genes as in MAG-1207, including an incomplete Wood-Ljungdahl pathway and the Rnf complex, though it lacks pyruvate formate-lyase and glycine reductase.

Neither of these Bipolaricaulota MAGs contains any known hydrogenases or formate dehydrogenases. A divergent FDH-like sequence that appears to be homologous with those in the *Methanosarcinaceae*, ANME-1, and *Thermodesulfovibrionales* MAGs was observed in multiple initial BinSanity bins classified as Bipolaricaulota (**Figure 6**, but it was not included in the three re-assembled, refined Bipolaricaulota MAGs. The most similar sequence in the NCBI NR database was from a MAG assembled by (Parks et al., 2021) from a Voltri Massif serpentinite spring (Brazelton et al., 2017).

A third Bipolaricaulota MAG (1503 CDS; 92% complete with 1% redundancy) was classified as genus UBA3574 within family Bipolaricaulaceae. It encodes a nearly complete Wood-Ljungdahl pathway, including the NADP-dependent formate dehydrogenase (FdhA) that is typical of acetogens. It does not encode the Rnf complex, but it contains genes encoding all subunits of the heterodisulfide reductase complex MvhAGD-HdrABC. It encodes a [NiFe] hydrogenase (HoxYH) and at least a partial gene cluster for membrane-bound hydrogenase, including the large catalytic subunit MbhL (**Figure 5; Supplemental Figure S8**). The predicted MbhL sequence is most closely related to two Bipolaricaulota MAGs from hydrothermal systems: the

Mid-Cayman Rise (Zhou et al., 2020) and Guaymas Basin (Dombrowski, Teske & Baker, 2018). Bipolaricaulota MAG-1503 also encodes an oxalate:formate antiporter distinct from those found in the *Methanosarcinaceae*, NPL-UPA2, and *Natrinincolaceae* MAGs (31-33% amino acid identities).

***Thermodesulfovibrionales* MAG**

Two ASVs classified as *Thermodesulfovibrionia* (a class within phylum Nitrospirae) differed from each other by a single base. One of these was the top ASV in each of the fluid samples from Markers 3 and C (8-24% of all sequences), while the second ASV dominated the Calypso fluids (7-17% of all DNA fractions and 27% of the RNA fraction). Lost City *Thermodesulfovibrionia* metagenomic sequences and ASVs were notably rare in the low-sulfide fluids from Camel Humps and the high-sulfide fluids from Marker 2 (**Figures 2-3**).

Accordingly, a MAG (1293 CDS; 92% complete with 0% redundancy) classified as order *Thermodesulfovibrionales* within class *Thermodesulfovibrionia* was most abundant in Marker 3 and Calypso fluids. For heterotrophic metabolism, it encodes a nearly complete glycolysis pathway and TCA cycle, plus genes for lactate dehydrogenase, pyruvate ferredoxin oxidoreductase (PorABCD), and an oxalate:formate antiporter distinct from those found in the *Methanosarcinaceae*, NPL-UPA2, and *Natrinincolaceae* MAGs (21-24% amino acid identities),

*Thermodesulfovibrionales* MAG-1293 has a partial Wood-Ljungdahl pathway and a monomeric CO dehydrogenase (CooS) with two maturation factors (CooF and CooC). It encodes the NAD(P)-dependent formate dehydrogenase typical of acetogens (FdhA), as well as aerobic formate dehydrogenase (FdoG) and the divergent FDH-like sequence also observed in other Lost City MAGs (**Figure 6**). It encodes a [NiFe]-hydrogenase (HyaAB) classified as [NiFe] Group 1c, a group of respiratory $H_2$-uptake hydrogenases that can use fumarate, sulfate, or metals as terminal electron acceptors. The MAG contains only one subunit of membrane-bound hydrogenase (MbhJ), which shares 43-49% amino acid identities with the MbhJ sequences of the other Lost City MAGs shown in **Figure 5**. The key genes required for nitrogen fixation (nifHDK) and dissimilatory sulfate reduction (dsrAB) are present.

***Desulfotomaculum* MAGs**

Two MAGs classified as family *Desulfotomaculaceae* were resolved based on their distinct coverage patterns. Both MAGs were estimated to be 94% complete with 0% redundancy, but one had more predicted genes (1580) than the other (1144). *Desulfotomaculaceae* MAG-1580 was only abundant in one sample from Sombrero, while MAG-1144 was abundant in Marker 3, Sombrero, and Calypso fluids (**Figure 3**). Several ASVs were classified as genus *Desulfotomaculum* within the family *Desulfotomaculaceae*. One of these was most abundant in Marker 3 (up to 14% of all sequences) and Calypso (2-8% of all sequences), roughly matching the distribution of MAG-1144, while another ASV (differing from the first by four bases) was only abundant in Sombrero fluids, similar to MAG-1580 (**Figure 2**).

Both *Desulfotomaculum* MAGs encode a complete or nearly complete glycolysis pathway and at least two genes of the TCA cycle (malate dehydrogenase and isocitrate dehydrogenase). In addition, *Desulfotomaculum* MAG-1144 has succinate dehydrogenase (SdhABC) and the beta subunit of fumarate dehydratase. *Desulfotomaculum* MAG-1580 encodes pyruvate ferredoxin oxidoreductase (PorABCD) and pyruvate formate-lyase (PflD). Both MAGs have a cation/acetate symporter (ActP), and MAG-1580 encodes a phosphonate transporter (PhnCDE).

Both MAGs have incomplete Wood-Ljungdahl pathways that lack the key enzyme CODH/ACS. Both MAGs also have monomeric CO dehydrogenase (CooS) and aerobic formate dehydrogenase (FdoG). No [NiFe] hydrogenases were detected, and only one subunit of [FeFe] hydrogenase (HndC) was present in the *Desulfotomaculum* MAGs. This hydrogenase is capable of $H_2$ oxidation with reduction of NADP in some organisms (Kpebe et al., 2018), but the presence of only one subunit in multiple Lost City MAGs (**Supplemental Table S5**) is curious and has unknown implications for the ability of these organisms to metabolize $H_2$.

The two *Desulfotomaculum* MAGs, in addition to the *Thermodesulfovibionales* MAG, are the only MAGs reported here that encode dissimilatory sulfite reductase (DsrAB). *Desulfotomaculum* dsrAB sequences were most abundant in Sombrero fluids, while *Thermodesulfovibrionales* dsrAB sequences were most abundant in Marker 3 and Calypso fluids. The two *Desulfotomaculum* MAGs also have the key genes required for nitrogen fixation (nifHDK).

#### ***Natronincolaceae* MAGs**

Two MAGs were classified as family *Natronincolaceae* within the Clostridia and shared 81-89% ANI with a MAG reconstructed from seafloor borehole fluids at North Pond (Tully et al., 2018). Other genera within the family *Natronincolaceae* include *Alkaliphilus* and *Serpentinicella*, which have been isolated from the Prony Bay hydrothermal field (Mei et al., 2016; Postec et al., 2021). The coverage of the *Natronincolaceae* MAGs was primarily in Sombrero fluids. The two MAGs shared 80% ANI but showed a few potentially important differences in their genomic inventories.

*Natronincolaceae* MAG-2163 (2163 CDS; 90% complete with 3% redundancy) has one of the largest genomes in this study, and it has an incomplete glycolysis pathway and incomplete TCA cycle. It encodes at least three steps of the Wood-Ljungdahl pathway, monomeric CODH (CooS), the Rnf complex, and the electron carriers HdrABC and MvhD.

In contrast, *Natronincolaceae* MAG-1138 (1138 CDS; 75% complete with 0% redundancy) has a smaller genome, a nearly complete glycolysis pathway, and no TCA cycle. It includes two genes associated with the Wood-Ljungdahl pathway, but not CODH, Rnf, Hdr, or MvhD. It does include acetate kinase (AckA) and phosphotransacetylase (Pta). It lacks any genes for ATP synthase, suggesting an obligate fermentative lifestyle.

*Natronincolaceae* MAG-2163 has [FeFe] hydrogenase (HndBCD), while MAG-1138 only has the HndC subunit. Both *Natronincolaceae* MAGs have at least one subunit of pyruvate dehydrogenase and pyruvate ferredoxin oxidoreductase, though they differ in which subunits they include (**Supplemental Table S5**).

***Dehalococcoidia* MAGs**

The most abundant *Dehalococcoidia* MAG (844 CDS; 73% complete with 0% redundancy) was classified as family SpSt-899 within order SZUA-161. It was one of the highest coverage MAGs in Calypso fluids as well as the Sombrero and Marker 2 metatranscriptomes. Unlike the other two *Dehalococcoidia* MAGs described below, glycolysis and the TCA cycle are incomplete, and there are no genes required for the degradation of large organic compounds. However, the presence of pyruvate:ferredoxin oxidoreductase, pyruvate formate-lyase, and an oxalate/formate antiporter suggest the ability to ferment low-molecular-weight organic compounds. In addition, the MAG has two steps of the Wood-Ljungdahl pathway (FchA and MetF), the Rnf complex, and a complete set of genes encoding the key enzyme CODH/ACS. No hydrogenases or formate dehydrogenases are present. Curiously, *Dehalococcoidia* MAG-844 has V(A)-type ATP synthase, which is typically associated with archaea, but has also been observed in Chloroflexi and Parcubacteria in another serpentinite-hosted spring (Suzuki et al., 2017).

Two additional *Dehalococcoidia* MAGs belong to the SAR202 cluster, one of which was classified by GTDB as order SAR202 and one as order UBA3495. Each MAG has 98% ANI with previously published marine Chloroflexi MAGs (**Supplemental Table S4**). *Dehalococcoidia* MAG-2669 (2669 CDS; 86% complete with 4% redundancy) has moderately high coverage in all chimney fluids except for Marker 3. *Dehalococcoidia* MAG-2875 (2875 CDS; 87% complete with 1% redundancy) was less abundant than MAG-2669 in all samples but otherwise exhibited a similar distribution pattern. Both MAGs encoded nearly complete glycolysis pathways and TCA cycles, and they have a variety of genes associated with the oxidation of various organic molecules, consistent with previous studies of marine Chloroflexi (Liu et al., 2021; Landry et al.). Both MAGs encode cytochrome c oxidase (CoxA), which represents the only evidence for aerobic respiration in any of the final, refined MAGs. *Dehalococcoidia* MAG-2875 has both (aerobic) pyruvate dehydrogenase and (anaerobic) pyruvate:ferredoxin oxidoreductase, as reported for other SAR202 genomes (Mehrshad et al., 2018), while MAG-2669 has neither. Neither MAG encodes any FDH or hydrogenases. MAG-2669 is predicted to encode sulfite reductase (Sir), adenylylsulfate reductase (AprAB), and sulfate adenylyltransferase (Sat), but dissimilatory sulfite reductase (DsrAB) is not present. Both MAGs have genes associated with the metabolism of organosulfur compounds, similar to those reported by (Mehrshad et al., 2018) for deep-sea SAR202 genomes (**Supplemental Table S5**).

All three *Dehalococcoidia* MAGs include genes for glycine reductase, thioredoxin, and selenocysteine synthesis. GrdB (beta subunit of glycine reductase) sequences from the two Lost City Dehalococcoidia MAGs that belong to the SAR202 marine cluster (MAG-2669 and MAG-2875) are distinct from the GrdB of *Dehalococcoidia* MAG-844 and from the GrdB of all other Lost City MAGs (**Supplemental Figure S11**).

### NPL-UPA2 MAGs

Three MAGs classified as candidate phylum NPL-UPA2 (new name Candidatus Horikoshi bacteria proposed by (Suzuki, Nealson & Ishii, 2018) were recovered, each only 55-72% complete and with distinct coverage patterns among the hydrothermal fluid samples. No ASVs or unclassified reads were classified as NPL-UPA2, as this group was not yet represented in the SILVA or Kaiju databases. NPL-UPA2 MAG-914 (914 CDS; 62% complete with 0% redundancy) and MAG-1083 (1083 CDS; 55% complete with 3% redundancy) were most abundant in Calypso fluids and nearly absent in all other locations. MAG-718 (718 CDS; 72% complete with 3% redundancy) was most abundant in Marker 3 fluids and exhibited a nearly inverse abundance distribution compared to the other two MAGs. All three NPL-UPA2 MAGs have incomplete Wood-Ljungdahl pathways that lack the first few steps and begin with methylene-THF dehydrogenase (FolD), and all three MAGs encode V(A)-type ATP synthase and the Rnf complex, as reported by (Suzuki, Nealson & Ishii, 2018).

In most other respects the two NPL-UPA2 MAGs that are prominent in Calypso fluids differ from the MAG that is most abundant in Marker 3 fluids. The Calypso MAGs encode multiple subunits of the CODH/ACS enzyme, while MAG-718 has only the methyltransferase subunit (AcsE). MAG-1083 also includes acetate kinase (AckA), phosphotransacetylase (Pta), and a cation/acetate symporter (ActP). In contrast, MAG-718 is the only NPL-UPA2 MAG that encodes pyruvate formate-lyase (PflD), the oxalate/formate antiporter, and carbonic anhydrase.

Evidence for hydrogenases in the NPL-UPA2 MAGs is lacking. Membrane-bound hydrogenase (MbhL) was identified in two of the MAGs by GhostKoala, but Prokka annotated these sequences as NAD(P)H-quinone oxidoreductase and formate hydrogenlyase. Neither sequence could be placed in the mbhL phylogeny of **Figure 5**. MAG-1083 includes one subunit of $F_{420}$-reducing hydrogenase (FrhB) and one subunit of [FeFe] hydrogenase (HndD), but the functional roles of these predicted proteins are unclear.

Unlike the NPL-UPA2 MAG reported by (Suzuki, Nealson & Ishii, 2018), none of the Lost City NPL-UPA2 MAGs have formate dehydrogenase nor the electron carriers Hdr and Etf.

**Patescibacteria MAGs**

Several MAGs classified as candidate phylum Patescibacteria were represented by two classes: Paceibacteria and Gracilibacteria (according to GTDB taxonomy). Paceibacteria MAGs (55-86% completion with 0-3% redundancy) had small genomes (as low as 307 kb with 63% estimated completion) with low GC content (24-45%). Gracilibacteria MAGs (61-83% completion with 0-1% redundancy) had even lower GC content (22-34%) and somewhat larger genomes but with very low annotation success. As few as 8% of all coding sequences in Gracilibacteria MAGs yielded GhostKoala results. Two of the Paceibacteria MAGs were most abundant in Marker 3 fluids, while Gracilibacteria MAGs were notably absent in Marker 3 fluids and were more abundant in Camel Humps and Sombrero fluids (**Figure 3; Supplemental Table S4**).

In general, Paceibacteria and Gracilibacteria MAGs included genes for the biosynthesis of key cellular components and basic information processing, but genes specific to catabolic pathways were rare. The few exceptions included a lactate transporter (LctP) that was present in the two highest-coverage Paceibacteria MAGs, and one of these MAGs also has lactate dehydrogenase (LdhA). In addition, acetate kinase (AckA), acylphosphatase (AcyP), and phosphoenolpyruvate synthase (PpsA) were present in one or more Paceibacteria MAGs (**Supplemental Table S5**). Propionyl-CoA carboxylase (PccB) was included in one Gracilibacteria MAG. All Paceibacteria MAGs, one of the Gracilibacteria MAGs, and almost all other MAGs in this study have a substrate-binding protein associated with the peptide/nickel transport system (K02035; **Supplemental Table S6**).

Curiously, methylene tetrahydrofolate dehydrogenase (FolD), which is part of the Wood-Ljungdahl pathway, was encoded by three Paceibacteria MAGs and one Gracilibacteria MAG. FolD was also observed in 17 MAGs from The Cedars, a serpentinite-hosted spring in California, that were classified as candidate phylum OD1 (Suzuki et al., 2017), now included within Paceibacteria in GTDB.

As mentioned in the main text, ATP synthase was completely absent in three of the Paceibacteria MAGs, and one Gracilibacteria MAG included only a single subunit. One Paceibacteria MAG encodes a V(A)-type ATP synthase instead of the F-type ATP synthase present in the other Paceibacteria and Gracilibacteria MAGs. MAGs that were classified as candidate phylum OD1 from The Cedars also encoded V(A)-type ATP synthase or lacked any ATP synthase at all (Suzuki et al., 2017).

**WOR-3 MAG**

One of the highest-coverage MAGs in Marker 3 and Sombrero fluids was classified as candidate phylum WOR-3, which was previously identified in methane-rich marine sediments (Baker et al., 2015) and was proposed to be renamed as Candidatus Stahlbacteria (Dombrowski et al., 2017) The WOR-3 MAG (59% completion with 0% redundancy) has a partial glycolysis pathway, perhaps explainable by the incompleteness of the MAG, and no TCA cycle. Other genes possibly indicative of organic carbon catabolism include those predicted to encode one subunit of pyruvate dehydrogenase (PdhD), pyruvate formate-lyase (PflD), formate dehydrogenase (FdoG), and a cation/acetate symporter (ActP). Like the NPL-UPA2 MAGs, one Paceibacteria MAG, and the archaeal MAGs, the WOR-3 MAG encodes a V(A)-type ATP synthase.

The WOR-3 formate dehydrogenase shares a maximum of only ~31% amino acid identities with any other sequences in the NCBI NR and JGI IMG "all isolates" databases. This divergent FDH-like sequence in the WOR-3 MAG is distinct from the divergent FDH-like sequence described above for other Lost City MAGs (**Figure 6**), and in both cases, additional research is required to establish whether their functions are indeed associated with formate metabolism.
